# Muscle spindles provide flexible sensory feedback for movement sequences

**DOI:** 10.1101/2024.09.13.612899

**Authors:** William P. Olson, Varun B. Chokshi, Jeong Jun Kim, Noah J. Cowan, Daniel H. O’Connor

**Affiliations:** The Solomon H. Snyder Department of Neuroscience, Kavli Neuroscience Discovery Institute, Zanvyl Krieger Mind/Brain Institute, Johns Hopkins University, Baltimore, MD 21218; Department of Mechanical Engineering, Johns Hopkins University, Baltimore, MD 21218

## Abstract

Sensory feedback is essential for motor performance and must adapt to task demands. Muscle spindle afferents (MSAs) are a major primary source of feedback about movement, and their responses are readily modulated online by gain-controller fusimotor neurons and other mechanisms. They are therefore a powerful site for implementing flexible sensorimotor control. We recorded from MSAs innervating the jaw musculature during performance of a directed lick sequence task. Jaw MSAs encoded complex jaw–tongue kinematics. However, kinematic encoding alone accounted for less than half of MSA spiking variability. MSA coding of kinematics changed based on sequence progression (beginning, middle, or end of the sequence, or reward consumption), suggesting that MSAs are flexibly tuned across the task. Dynamic control of incoming feedback signals from MSAs may be a strategy for adaptable sensorimotor control during performance of complex behaviors.

---

Complex behavior is built out of simpler motor actions. Past work has shown how hierarchically structured circuits can enact motor control at multiple levels (i.e. by driving single actions, or by organizing behavior across actions)^1–3^. Motor control is implemented through sensorimotor loops that continuously integrate sensory feedback into ongoing motor commands. The relevance of sensory feedback, however, can change based on task demands^4–6^. It is unclear if feedback dynamics during complex behavior may reflect multiple levels of hierarchical control.

Muscle spindles, innervated by sensory muscle spindle afferents (MSAs) and by gain-controller efferent fusimotor neurons, are an important source of feedback about body position and movement. MSAs show complex and variable encoding of kinematics. During stretch of passive muscles, MSA responses are correlated with muscle length and its time derivatives^7^. However, during active movement, MSA activity can be dramatically decoupled from body kinematics^8–10^. The role of this complex activity has been debated^11,12^. Recent models^13^ have indicated that MSAs may provide flexible movement representations that can adapt to task requirements.

Here we recorded from MSAs during execution of flexible, learned motor sequences. We used a recently developed task in which head-fixed mice perform complex sequences of directed licks to receive a water reward^3^. Recordings from MSA neurons innervating the jaw (located in the mesencephalic trigeminal nucleus, abbreviated as MEV for the “mesencephalic nucleus of the fifth (V) cranial nerve”) revealed complex and diverse task-related responses. A component of the MSA activity could be explained by jaw and tongue kinematics. However, much of the activity was instead linked to sequence- and task-level variables rather than kinematics, likely due to online modulation of MSA responses. Our results indicate that motor systems may use dynamic tuning of first-order sensors (MSAs) to strategically shape incoming sensory feedback for hierarchical motor control.

## Results

We trained head-fixed mice to perform the directed lick sequence task, as described previously^3^. The task structure features licks across orofacial kinematic space in two sequence directions, as well as post-reward ‘drinking’ licks at the end of each trial. Mice licked a moving port to drive it through an arc of seven locations surrounding the mouth (Figure 1a) after an auditory cue. Sequences started on either the left or the right and progressed to the opposite side, and each trial terminated with delivery of a water reward through the port and a series of consummatory licks. The next trial started with a sequence in the opposite direction after a random intertrial interval. We tracked jaw and tongue kinematics from high-speed (400 Hz) video of the animal’s mouth, extracting 3 jaw parameters (mandible tip positions: dorsoventral *z*, mediolateral *x*, anteroposterior *y*), 3 tongue parameters (tongue length *L*, angle from the midline *θ*, angle from the vertical *φ*), and the associated velocities (first derivatives *ż*, *ẋ*, *ẏ*, *L̇*, *θ̇*, *φ)̇* (Figure 1b). We defined two additional task variables: lick cycle phase, the relative position of the jaw within a single lick cycle, and ‘progress’, the fraction of the trial that has been completed (relative to water delivery; Figure 1d–e; Methods).

**Figure 1.**
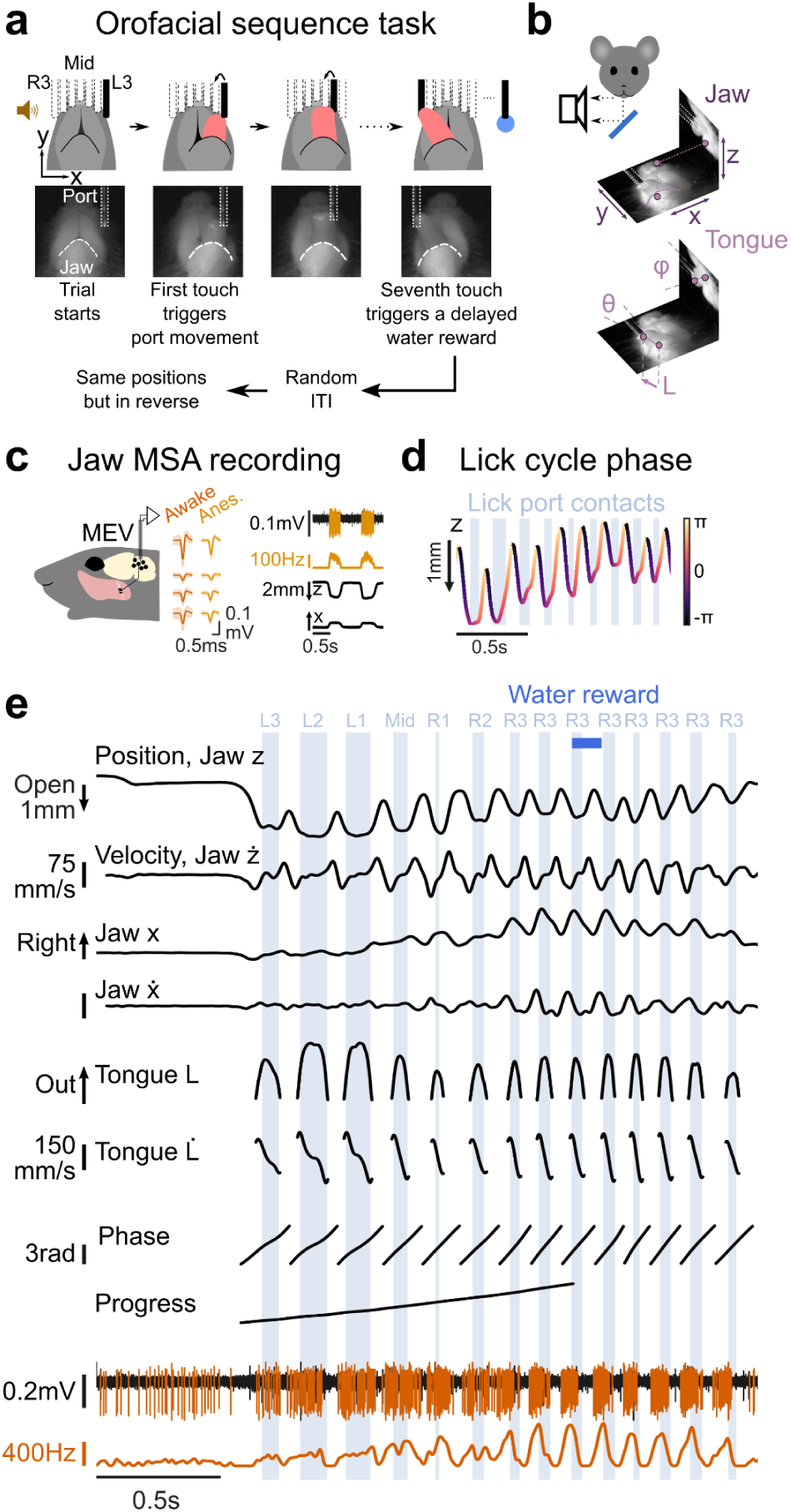
Jaw MSAs were recorded during performance of an orofacial sensorimotor sequence. **a,** Schematic of the sequence licking task. **b,** 6 jaw and tongue kinematic parameters were extracted from dual-view high speed (400 Hz) video of the mouth. 3 jaw parameters (*z*, *x*, *y*) described the distal jaw position, and 3 tongue parameters (*L*, *θ*, *φ*) described the tongue position. **c,** Left, we isolated single jaw MSAs from tetrode recordings in the midbrain mesencephalic nucleus of head-fixed mice. Middle, waveforms (mean ± std. dev.) of a representative jaw MSA in awake and anesthetized conditions. Right, unit responses to passive jaw movement under anesthesia, showing the selected channel extracellular recording and smoothed (7.5 ms std. dev. Gaussian kernel) spike rate (top) as well as simultaneous jaw kinematics (bottom). **d,** Representative lick cycle phase assignments, extracted from jaw *z* position data, for 9 licks. **e,** Sample trial time-series data, showing extracted kinematics (*z*, *x*, *L* and velocities *ż*, *ẋ*, and *L̇*), phase, progress, selected channel extracellular recording, and unit smoothed spike rate.

MSA neurons innervating the jaw musculature were isolated from chronic 32-channel tetrode recordings in MEV (n=78 unique units from 7 animals) (Figure 1c; Supplemental Figure 1) during sequence task performance (Figure 1e). We searched for the muscles innervated by MSAs by gently probing the cheek musculature under isoflurane anesthesia (Supplemental Figure 2) and identified cheek locations for some units (17/78). We found five characteristic response types. Four of these types were associated with silent baseline activity and could be localized to specific regions of the jaw musculature^14^: *temporalis*, *posterior masseter*, *anterior masseter*, and *ventral masseter – silent baseline*. A fifth type comprised units showing tonic activity under anesthesia with silencing/activating response fields localized to the ventral masseter (*ventral masseter – tonic baseline*). This last type may innervate the pterygoid muscles sitting inside the mandible (Discussion).

### Lick- and sequence-level task dynamics drove MSA responses

We aligned kinematics and MSA spike rates across trials using lick-cycle phase (Figure 2a–b). Jaw MSAs were strongly modulated during task performance, with highly diverse responses across the population. Many MSAs showed strong modulation on the order of single lick events (phase tuning), while others showed weaker response modulation to individual licks (‘slowly modulated’ units). We segregated units on the basis of the amount of mutual information^15^ single-unit spiking held with lick-cycle phase (Methods) and identified 47/78 phase coupled units (Figure 2c).

**Figure 2.**
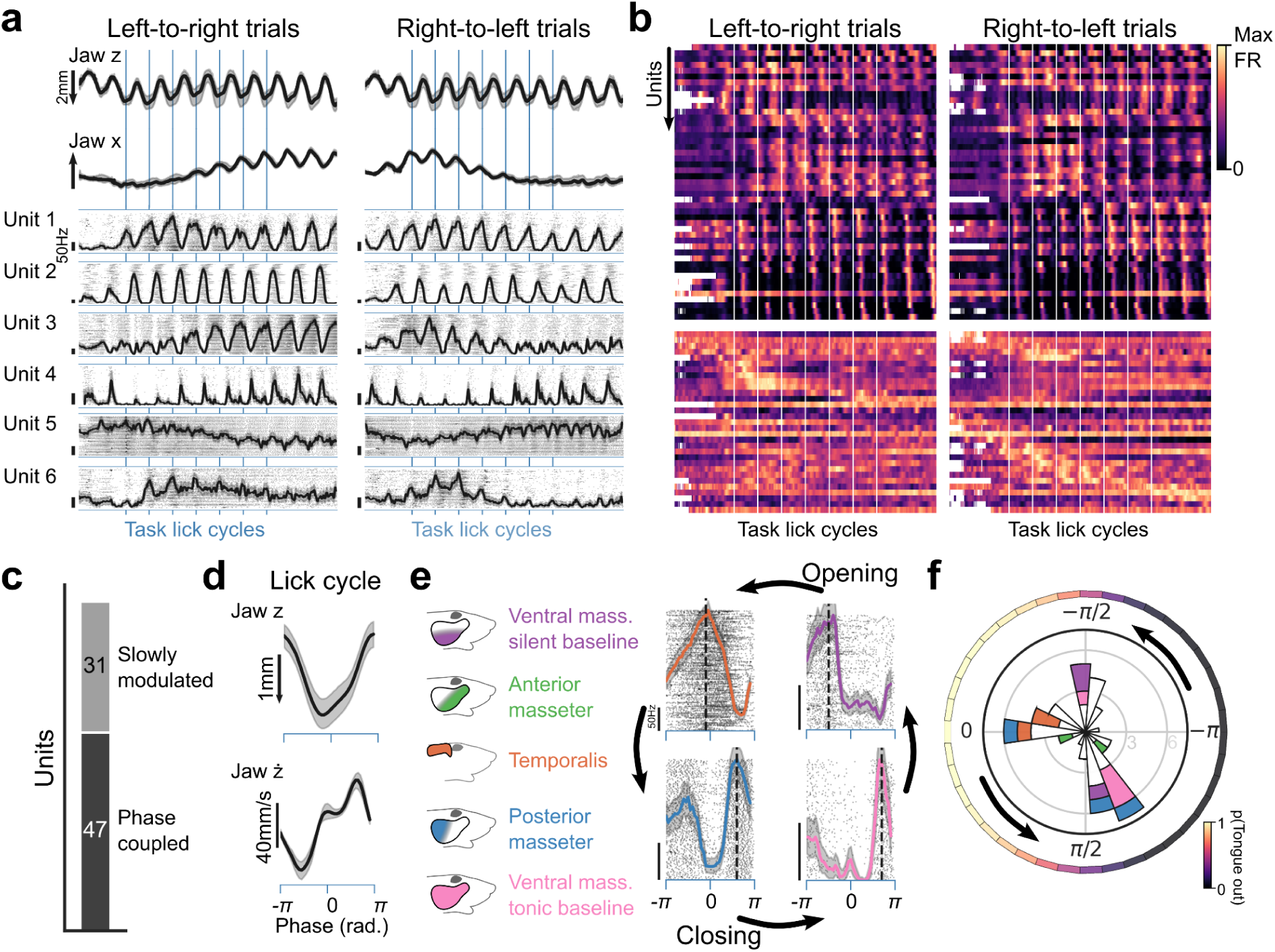
Lick- and sequence-level modulation of jaw MSA activity. **a,** Phase-aligned trial (−1.5 to +9.5 cycles from first contact lick) averaged kinematics (across animal and session mean ± bootstrapped std.) and unit rasters with histograms (± bootstrapped std.) for six units and two sequence directions. **b,** Heatmaps of phase-aligned unit histograms (normalized to maximum rate) for two sequence directions. Top, phase coupled units, ordered by phase tuning peaks. Bottom, non-phase coupled (i.e. ‘slowly modulated’) units, ordered by position of maximum firing across the two sequences. **c,** Numbers of phase coupled and non-phase coupled units (Methods). **d,** Phase-averaged kinematics (across animal and session mean ± bootstrapped std.). **e,** Phase-aligned unit rasters and histograms (± bootstrapped std.) for four units. Dashed lines indicate primary phase tuning peaks, color indicates cheek probe response field. **f,** MSA unit phase tuning. Inner, primary phase tuning peak polar histogram (bin size = *π*/10 rad.). Colors indicate cheek probe response field. Unit phase tuning was correlated with peripheral innervation site. Outer, tongue out probability based on lick cycle phase (Methods).

The musculature of the jaw is highly asymmetric, with large muscles driving closure and comparatively small, weak muscles driving opening^14^. One might expect a similar asymmetry among the jaw MSA phase responses, with most MSAs activated by jaw muscle stretch during the opening swing. In contrast to this prediction, we identified phase tuning peaks for the phase coupled subset of MSA units and observed tiling of responses across the lick cycle (including during closing phases) (Figure 2d–f). MSAs fired either during the opening swing (just prior to tongue protrusion), around the time of maximal opening, or during the closing swing (Figure 2f). Phase tuning is likely determined by the muscle of innervation, and units generally segregated based on their peripheral response fields. For example, while *temporalis* units fired around maximal opening, *ventral masseter – tonic baseline* units fired during the closing swing (Figure 2f).

MSA phase tuning may be driven by stretch sensitivities and/or top-down modulatory drives during lick events. Some MSAs fire during muscle contraction due to parallel activation of α motor and fusimotor neurons^9^. We asked if phase tuned activity of some MSAs may co-occur with jaw muscle contraction. We recorded *temporalis* and *masseter* muscle activity (EMG; Methods) in a separate cohort of animals (n=16 sites from 4 animals) performing the sequence licking task (Supplemental Figure 4a-b). Jaw muscle activity peaked at two points of the lick cycle: once during the opening swing around the time of tongue protrusion, and once during closing (Supplemental Figure 4c-d). The first peak may stabilize the jaw for tongue protrusion, while the second likely drives jaw closure. Based on these results, the phase tuning of some MSAs (i.e. tuning to closing phases) may represent contraction-linked MSA activity.

While some MSAs responded consistently throughout the motor sequences (Figure 2a, Units 1 and 2), most showed dynamic modulation of their response across the sequences. This included both phase coupled and slowly modulated units (Figure 2b). Notably, we observed responses linked to kinematic changes over the sequence (i.e. tuning to leftward or rightward licks) (Figure 2a, Units 3 and 5) as well as responses linked to progress regardless of sequence direction (Figure 2a, Units 4 and 6). Activity across the population tiled the sequences, and many units showed stronger activity in one of the two sequence directions (Figure 2b). Together, these recordings indicate diverse encoding of kinematics across the population, and additionally indicate that MSAs may be driven by task dynamics that are independent from the execution of specific movements.

### Jaw–tongue kinematics partially explained jaw MSA activity

We observed diverse responses among jaw MSAs during sequence task performance. To ask if this activity might encode orofacial movements, we built encoding models that used the kinematics to predict single-unit activity. We compared the performance of these models across the population.

Jaw and tongue movement during the task was highly coordinated. However, jaw and tongue kinematics were only partially correlated (Supplemental Figure 5a). We identified kinematic coordinate axes from jaw, tongue, or jaw+tongue time-series data using Singular Value Decomposition (SVD) (Figure 3a–b; Supplemental Figure 5c; Methods). Analysis of the cumulative variance explained based on the number of included axes (Figure 3c) reveals a high dimensionality to the orofacial movement space (∼8 dimensions are needed to explain >80% of jaw+tongue movement variability).

**Figure 3.**
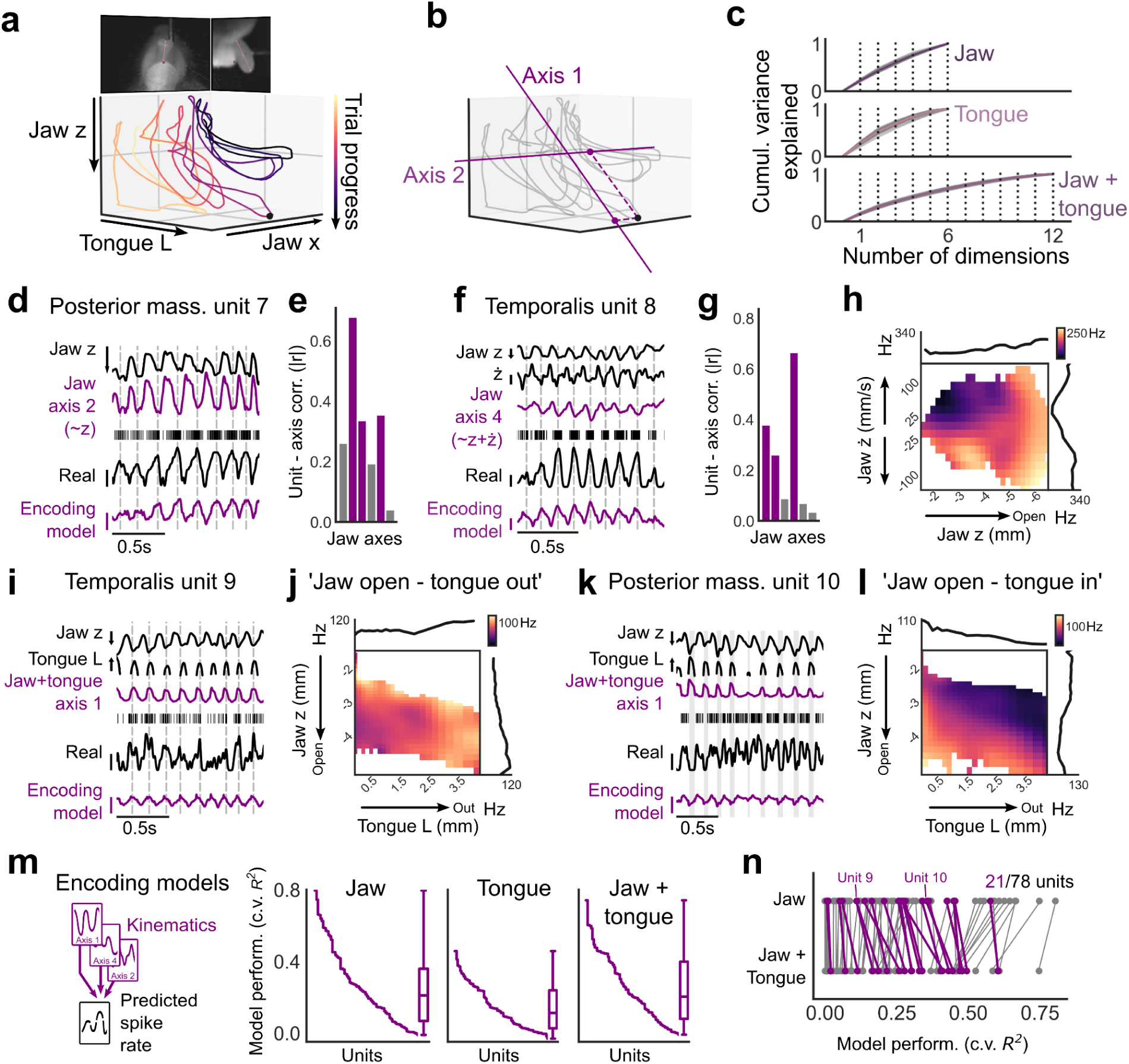
Jaw MSA encoding of complex tongue-jaw kinematics. **a,** Jaw *z*, jaw *x*, and tongue *L* trajectories during a sample trial. Black dot indicates the time point shown in the video frame. **b,** Extraction of coordinate axes from 3D (*z*,*x*,*L*) sample data, and projection of the chosen time point onto these axes. **c,** Cumulative variance explained based on the number of included coordinate axes calculated from 6D (positions and velocities) jaw, 6D tongue, or 12D jaw+tongue data. Gray lines indicate single sessions (n=31 sessions from 6 animals). Colored lines indicate across session mean. **d,** A unit innervating *posterior masseter* (unit 7) was highly correlated with a jaw coordinate axis that described instantaneous jaw opening position (∼*z*). Shown are the jaw *z* and jaw axis 2 weight trajectories along with unit spike rasters, unit smoothed (7.5 ms std. Gaussian kernel) spike rate, and linear encoding model predicted spike rate for selected licks. Dotted vertical lines mark peak unit spike rate within each lick cycle. **e,** Correlations of unit 7 spike rate and jaw coordinate axes. Purple bars indicate top 3 axes used in linear encoding models. **f*–*g,** Unit 8, innervating *temporalis* muscle, was highly correlated with a jaw coordinate axis that described jaw open position and velocity. **h,** Unit 8 co-tuning to jaw open position (*z*) and velocity (*ż*). Top, right, 1D *z* and *ż* tuning curves. Center, 2D tuning surface. The unit fired during downward opening movement. **i,** Activity of unit 9, innervating *temporalis* muscle, was correlated with a jaw+tongue axis that described jaw open, tongue protrusion movements. **j,** Unit 9 was co-tuned to jaw *z* and tongue *L*. **k***–***l,** Unit 10, innervating posterior masseter, was tuned to jaw opening, but silent during tongue protrusions. Gray bars in k indicate tongue protrusion out of the mouth. **m,** Mean Monte-Carlo cross-validated performance (*R*^2^) of linear encoding models, each of which predicted spike rates using weights onto the top 3 coordinate axes for each unit. Performance varied across MSAs, with relatively few units showing high encoder performance. **n,** 21/78 units (purple) showed significant outperformance of jaw+tongue vs jaw linear encoding models (Methods). Scale bars: *z*, 1 mm open, *ż*, 40 mm/s, *L*, 2 mm out, real and predicted unit activity, 100 Hz (d,i,k) or 200 Hz (f).

For our encoding models, we projected the time-series data onto the coordinate axes, calculated the weights, and then linearly regressed smoothed (Supplemental Figure 5b; Methods) single-MSA spike rates onto these weights. For each neuron, we chose the top 3 jaw, tongue, or jaw+tongue axes on the basis of correlation (absolute value of the Pearson correlation) with the smoothed spike rates. For a minority of units, the linear encoding models strongly predicted MSA activity (23/78 units have mean cross-validated *R*^2^ values > 0.5 for jaw+tongue models) (Figure 3m). However, major components of the MSA population activity were poorly explained by the linear encoding models. We additionally fitted more flexible decision-tree based models (Boosted trees^16^) to predict MSA activity using either the coordinate axes calculated from jaw–tongue kinematics or using features extracted via SVD of the video data itself (Facemap^17,18^) (Supplemental Figure 6). These models outperformed the linear encoding models, but left major components of the neural activity unexplained by kinematics (31/78 and 34/78 units with mean c.v. *R*^2^ > 0.5 for models fitted on jaw+tongue axes and video features, respectively).

We examined relationships between the activity of single MSAs and kinematics to understand the tuning of these neurons during behavior. Classic work has shown that MSAs encode combinations of the length of their resident muscle and its rate-of-change (velocity) under conditions of passive muscle stretch^7^. Indeed, we identified individual neurons whose activity was well explained by jaw position (Figure 3d–e) or by co-tuning to jaw position and velocity (Figure 3f–h), consistent with classic models.

We next asked if any MSAs had activity better explained by coordinated jaw+tongue kinematics compared to jaw kinematics alone. 21/78 units show significant outperformance of encoders using 3 jaw+tongue axes compared to 3 jaw axes (95% CI of jaw+tongue encoder - jaw encoder c.v. *R*^2^ > 0, difference taken within each resample for 1000 bootstrap resamples) (Figure 3n). One subset of these units showed tuning to jaw opening and tongue protrusion conformations (Figure 3i–j; Supplemental Figure 7a–c). A second subset was tuned to jaw opening but was silent during tongue protrusions (Figure 3k–l; Supplemental Figure 7a–c). Jaw–tongue tuning subtypes were related to peripheral response field locations, with ‘jaw open – tongue out’ responses seen in *temporalis* and *anterior masseter* units, and ‘jaw open – tongue in’ responses seen in *posterior/ventral masseter* units (Supplemental Figure 7c). These responses to coordinated jaw and tongue movements may reflect complex local changes in muscle stretch and/or changes in top-down modulation of MSAs. Notably, while unit kinematic co-tuning overlaps with orofacial conformations during lick port contacts, it was maintained in licks with no port contact (Supplemental Figure 7d–e). This indicates that these responses were not likely to be driven by external contact forces.

Overall, we found encoding of complex orofacial kinematics in jaw MSAs. While a small subset of MSAs strongly encoded movement (Figure 3m), large components of the responses across the population of MSAs were poorly explained by kinematics alone.

### MSAs were tuned to order within the sequence

During performance of the lick sequences, phase and progress represent ‘task variables’ that were linearly independent from the kinematics (Supplemental Figure 8). Based on our observation of MSA responses linked to sequence progress (Figure 2a, Units 4 and 6), we asked if these neurons may be specifically tuned to lick order independent of the kinematics.

We considered licks to the two outermost port positions (L3 and R3) obtained from sequences of both directions. These represent licks under two order conditions (‘first’ and ‘last’) at both locations (Figure 4a,c). We then sub-selected the licks that were best matched on the kinematics within each position (i.e. ‘L3 first’ vs. ‘L3 last’) (Supplemental Figure 9b,e,h), selecting equal numbers from each position–order combination (n=40 total licks). We excluded one session in which the lick kinematics could distinguish lick order (indicating insufficient matching of lick kinematics) (Supplemental Figure 9j). For each unit, we determined whether spiking during the licks could be used to classify lick position, order, or both. We pooled licks by position, or by order (Figure 4c), and built linear classifiers using lick mean spike counts. We performed Receiver Operating Characteristic (ROC) analysis on these classifiers, evaluating the performance using the ROC Area Under the Curve (AUC, 0.5 indicates chance level performance) (Figure 4d; Supplemental Figure 9c,f,i). Units responsive to position regardless of lick order (Unit 5 in Figures 2 and 4b,d; Supplemental Figure 9a–c) had strong position classifier performance and weak order classifier performance. Units that instead showed consistent activity based on sequence progress, regardless of sequence direction (Unit 11 in Figure 4b,d; Supplemental Figure 9d–f), had strong order classifier performance and weak position classifier performance. Some units could be used to classify both position and order (Supplemental Figure 9g–i), indicating tuning to position in a manner that depended on sequence direction. This could be related to sensitivity to the direction of movement, or to differences in top-down modulation between sequences.

**Figure 4.**
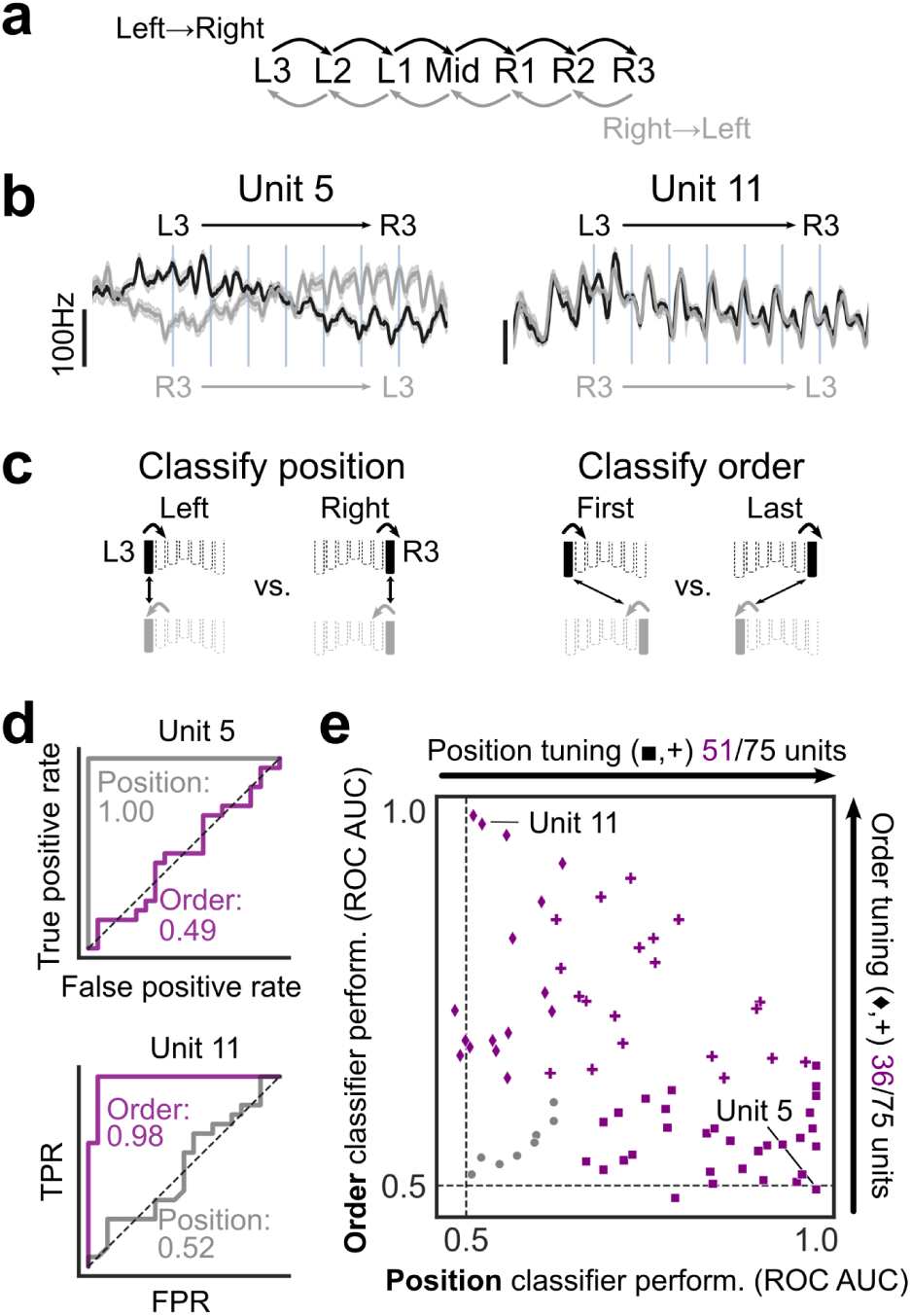
MSA task activity was driven by kinematics and by sequence progress. **a,** Task schematic for left-to-right and right-to-left sequences. **b,** Phase-aligned across-trial histograms for a unit tuned to lick port position (Unit 5, Figure 2) and a unit tuned to lick order (Unit 11). **c,** Schematic of position and order classifiers. Data were limited to task licks to the outer port locations (L3 and R3), and grouped either by position (left vs. right) or sequence order (first vs. last) (Methods). **d,** Receiver operating characteristic (ROC) analysis of classifiers that used unit mean spike counts during licks to classify position or order. Performance was measured as the area under the receiver operating curve (ROC AUC, 0.5 represents chance performance). **e,** Order vs. position classifier performance. Purple indicates significant performance (Methods). ◼, position-only significant. ♦, order-only significant. **+**, position and order significant.

Across the population, most units (67/75) showed significantly better than chance classification performance for position (31/75), order (16/75), or both (20/75) (95% CI of AUC > 0.5, estimated via bootstrap resampling on the licks for 1000 resamples) (Figure 4e). Position and order tuning was weakly related to unit peripheral response field (Supplemental Figure 9k). Repeating this analysis but using licks to the two positions on either side of the middle lick (L1 and R1) similarly showed order tuning of some MSAs (position-only: 28/60; order-only: 6/60; both: 12/60) (Supplemental Figure 9l). In summary, we identified units tuned to a task variable that is independent from the kinematics (lick order with the sequence), as well as units tuned to kinematics in a sequence-dependent manner.

MSA tuning to lick order is likely explained by task-driven changes in top-down modulatory drives. We asked if such task-driven changes might be seen in jaw muscle activity during sequence performance. While EMG activity is expected to be largely correlated with movements, it can be decoupled from body kinematics (for example, by co-contraction of antagonist muscles^19^). Jaw muscle EMG recordings showed diverse sequence responses, likely based upon the specific motor units sampled (Supplemental Figure 10a). We built linear classifiers that used activity from single recording sites to decode position or order of licks to the outermost positions (Supplemental Figure 10b-c). Similar to the MSAs, we found components of muscle activity linked to lick port position as well as lick order independent of position (position-only: 7/16 recordings; order-only: 2/16; both: 7/16). Therefore, task-driven changes in MSA responses (Figure 4, Supplemental Figure 9) co-occur with changes in output motor drives across the movement sequences (Supplemental Figure 10h).

### Sequence task and drinking licks differentially activated jaw MSAs

The sequence task featured licks that were directed to a series of specific port locations and drove movement of the port (‘drive’ licks), followed by a series of ∼5–10 water consummatory licks to a stationary port after reward delivery (‘drink’ licks) (Figure 5a). Sensorimotor cortex differentially encodes these two movement contexts, with stronger encoding of drive compared to drink licks in premotor cortex^3^.

**Figure 5.**
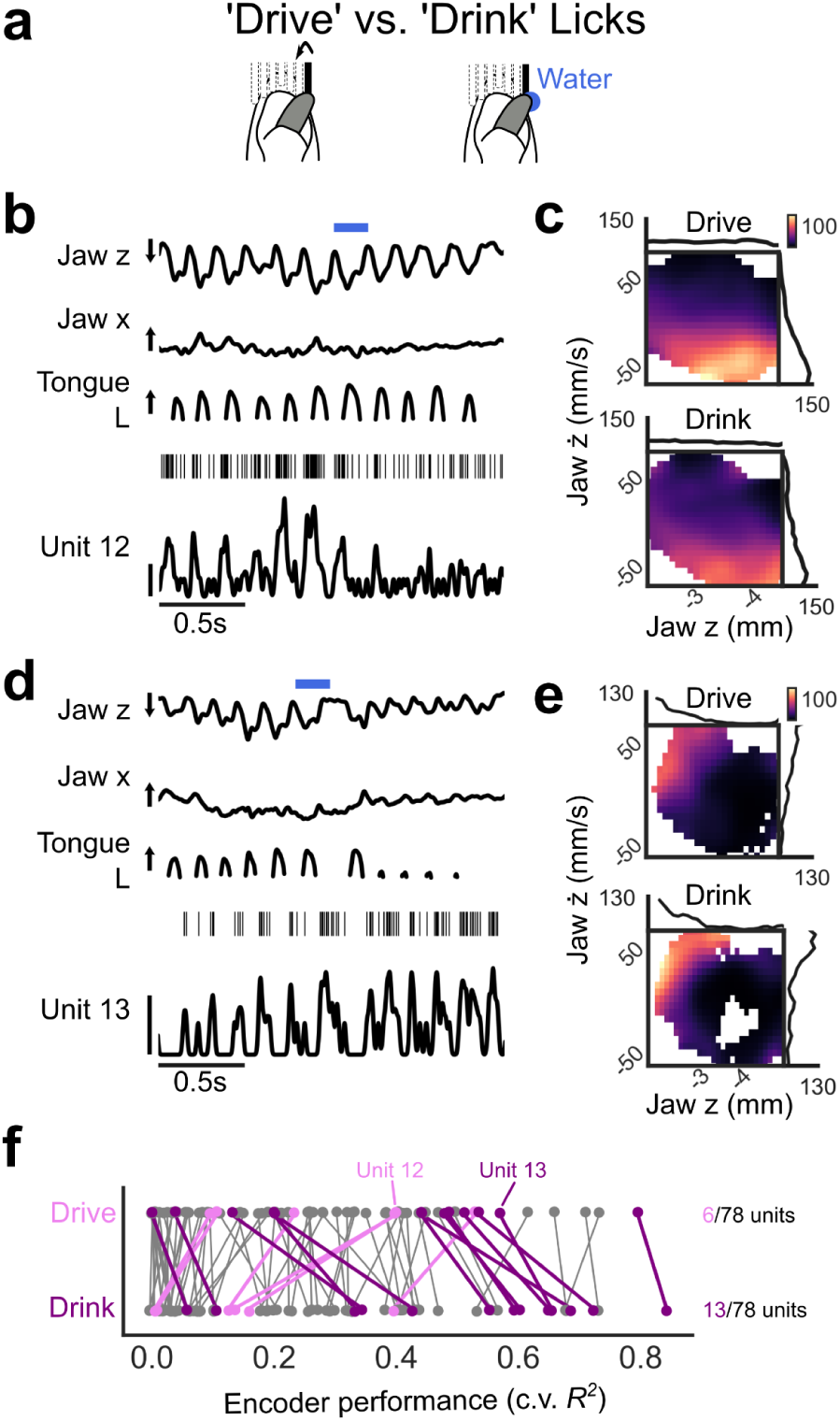
Jaw MSA activity reflected the task context of licks. **a,** ‘Drive’ licks were required to advance the port during the sequence, while ‘drink’ licks occurred after reward delivery. **b,** Kinematics, unit 12 spike rasters, and smoothed (7.5 ms std. Gaussian kernel) spike rates for licks before and after reward delivery. Unit 12 showed stronger coupling to drive licks. **c,** Unit 12 jaw *z* and *ż* tuning curves and 2D tuning surfaces for drive and drink licks. **d*–*e,** Unit 13 showed stronger coupling to drink licks. **f,** Performance of encoder models (top 3 jaw axes) fitted after sub-selecting data to match the drive and drink jaw movement spaces (Methods). 19/78 units showed a significant (Methods) difference in performance. Scale bars: *z*, 1 mm open, *x*, 1 mm right, *L*, 2 mm out, unit activity, 100 Hz.

Some MSAs responded differentially to licks before and after reward delivery, even when comparing licks with similar kinematics. We observed units with stronger responses to drive licks (Figure 5b), as well as units with stronger responses to drink licks (Figure 5d). We therefore asked if MSA activity was more strongly coupled to kinematics in one of these two movement contexts. Because drive licks explore a larger movement space than drink licks, we first limited data to regions of the kinematic space well represented by drink licks (Methods). 2D tuning histograms of jaw position and velocity showed dramatic changes in MSA kinematic tuning between contexts (Figure 5c,e).

To investigate this further, we fitted encoding models to predict MSA activity using jaw kinematics during drive or drink licks. 19/78 units showed a significant change in the strength of coupling to kinematics between conditions (95% CI of drink encoder performance - drive encoder performance did not include 0, comparison performed within resample for 1000 bootstrap resamples) (Figure 5f). This included units with stronger coupling to drive licks (6/78 units) as well as units with stronger coupling to drink licks (13/78 units). Drive and drinks licks showed different muscle activity patterns for some EMG recordings, with more complex and variable activity seen during drive licks (Supplemental Figure 10d-g). The shift in task context between drive and drink licks is likely associated with a change in sensorimotor control regimes.

### Jaw MSA ensembles encoded kinematic and task parameters

We observed MSA tuning to kinematics (Figure 3) as well as tuning to task parameters that are independent from the kinematics (Figures 4 and 5). We therefore asked if jaw MSAs show mixed encoding of kinematic and task information. We fitted linear models that use spike rates from simultaneously recorded jaw MSAs (‘ensemble decoding models’) (n=2–10 units per session) to predict kinematic and task parameters.

Jaw and jaw+tongue coordinate axis weights could be decoded from MSA activity (Figure 6a). We used an iterative dropout strategy to define the distribution of decodability across these ensembles (Methods). Removal of 1–2 top-performer units caused dramatic loss of decoder performance for many MSA ensembles (Figure 6b). This suggests that kinematic information is sparsely encoded among the ensembles.

**Figure 6.**
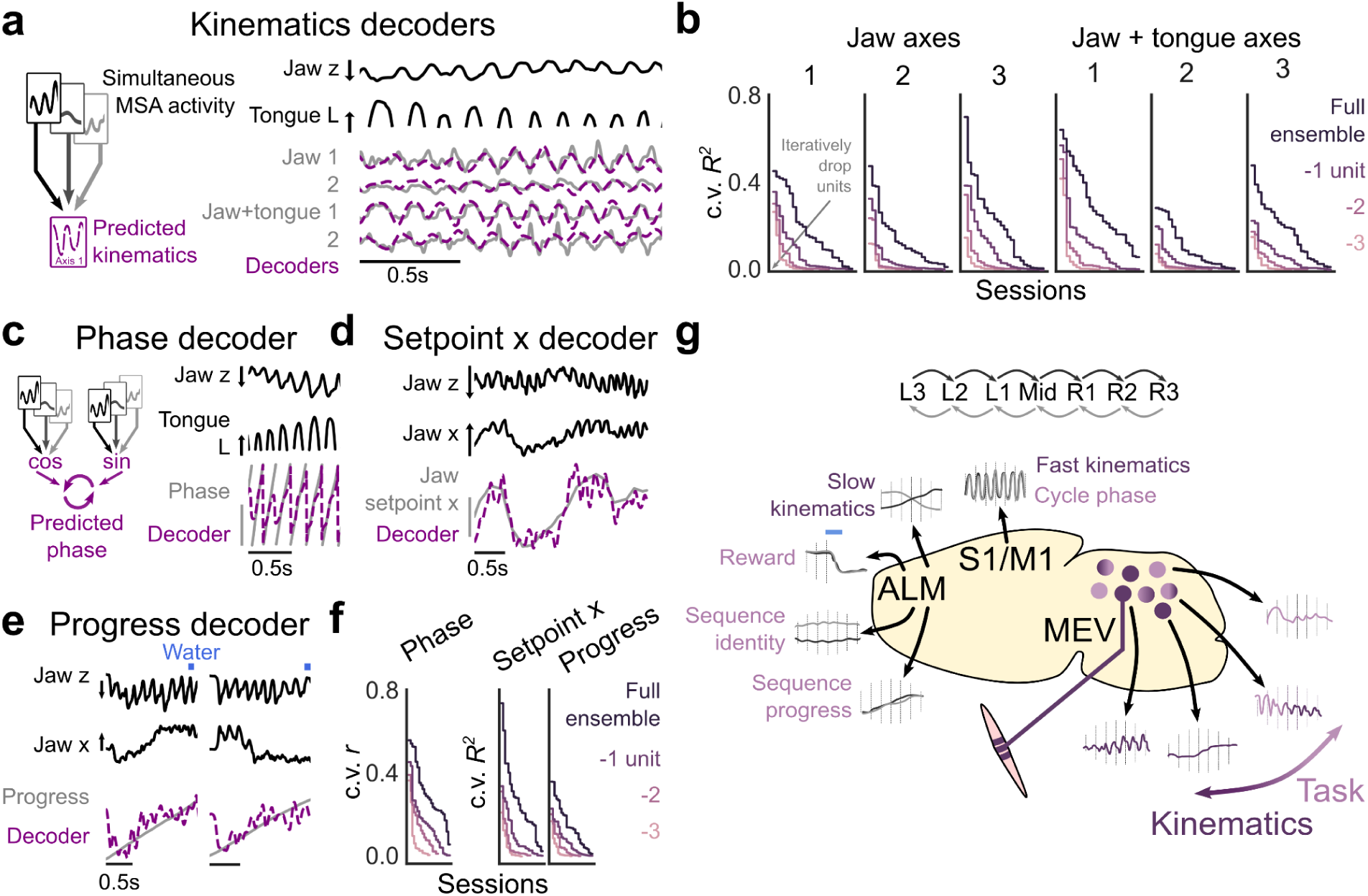
Mixed encoding of kinematics and task variables in MSA ensembles. **a,** Linear ensemble decoders predicted kinematics using simultaneously-recorded (2*–*10 units per session) unit spike rates. Right, representative time series data and decoder predictions from one session. **b,** Session kinematic decoder performances (5-fold cross-validated *R*^2^) for full ensembles and nested ensembles missing 1, 2, or 3 top-performer units. **c,** Decoding of lick cycle phase using linear models. Right, representative time series data and decoder prediction for one session. **d*–*e,** Linear decoding of jaw *setpoint x* (Methods) and progress. **f,** Session task variable decoder performances (5-fold cross validated Pearson’s *r* or *R*^2^) for full and nested ensembles. **g,** Sensorimotor cortex hierarchically encoded cycle-by-cycle and sequence-level information during this task^3^. We found a mixed representation of kinematic and sequence-task information in jaw MSAs. Scale bars: *z*, 2 mm (a,c) or 1 mm (d,e) open, *L*, 3 mm out, phase, 3 rad., *x*, 1 mm right, *setpoint x*, 1 mm.

Given the population tiling of responses across phases of the lick cycle, we asked if jaw MSAs could be used to decode lick cycle phase. To decode phase (a circular variable) using linear models, we mapped phase to 2D Cartesian coordinates (cos(*phase*), sin(*phase*)) and linearly decoded these components using neural activity. The predicted terms were then converted back to phase (Figure 6c). Jaw ensembles could be used to decode lick cycle phase (Figure 6c), with performance relying strongly on high-information units (Figure 6f).

Finally, consistent with our observation of lick position and order tuning (Figure 4), we found encoding of two parameters that together describe sequence performance: the jaw *setpoint x* position (representing the slow change in mediolateral center point of each lick) (Methods) and sequence progress (Figure 6d–f). Again, ensemble performance relied strongly on high-information units (Figure 6f).

Taken together, we found encoding of ‘fast’ (i.e. cycle-by-cycle) and ‘slow’ (i.e. sequence-level) kinematics as well as task variables that are independent of the kinematics. This information was sparsely encoded among the ensembles, indicating diverse mixed representations of kinematic and task variables in MSA populations (Figure 6g).

## Discussion

### Kinematic encoding by MSAs

MSAs comprise two major neuronal subtypes (primary, or type Ia, and secondary, or type II). Under passive stretch conditions (such as in the anesthetized animal), primary MSAs are co-tuned to muscle length and velocity, while secondary MSAs predominantly encode length^7^. MSAs report the direction of passively applied^20^ or actively generated^21^ movements. However, during active movement, MSA activity is only partially coupled to kinematics^9,10^. MSAs are mechanosensors driven by intra-spindle tension and its derivatives^8,22^. Much of the complex activity seen during active movement can be understood in terms of the impact of fusimotor (γ and β motor neuron) drive on spindle tensions via activation of the spindle intrafusal muscle fibers^8^. Fusimotor neurons are co-driven with the α motor neurons that cause muscle contraction (‘α-γ coactivation’)^11^, however they can also be independently controlled^13^. MSA outputs are also modulated via axo-axonic synaptic contacts onto their central arbors^23–25^, and the jaw MSAs additionally receive axo-somatic inputs onto their cell bodies in the midbrain MEV^26–29^.

We report that kinematic encoding models accounted for less than half of the observed MSA activity during active movement. While MSA coding can be highly nonlinear^7^, nonlinear models showed only moderate increase in performance compared to linear models (Supplemental Figure 6). Comparison across MSAs revealed a wide-distribution of kinematic encoding, with only a minority of units strongly coupled to the kinematics (Figure 3m). This included units primarily tuned to jaw position (Figure 3d) and units co-tuned to position and velocity (Figure 3f). Kinematic variables could be decoded from MSA ensemble activity in a manner consistent with sparse population coding (Figure 6a–b). It is possible that a minority of MSAs are well-positioned to stably encode specific kinematic parameters based on their location and/or response properties. We cannot account for movement hidden in our video data (for example, movement of the tongue within the mouth), which may drive some of the unexplained neural activity. Moreover, our observation of MSAs co-tuned to jaw and tongue kinematics (Figure 3; Supplemental Figure 7) suggests that higher-order circuits could use MSAs to control movement in an abstracted, multi-effector space.

### Peripheral topography and MSA phase tuning

The jaw muscles and bony structures are irregularly shaped. Most muscles lack tendons and instead attach to the complex structures of the skull and mandible with long aponeuroses^14^. We identified five characteristic fields of responsiveness to gentle cheek probing under anesthesia (Supplemental Figure 2). Four of these could be localized to regions of the largest jaw muscles: the *temporalis* and the *masseters* (comprising 2 muscles in the mouse, the deep and superficial masseters). A fifth field (‘*ventral masseter* – *tonic baseline*’) may represent units innervating the internal pterygoid, which sits inside of the mandible and elevates the jaw^30^. The slight jaw depression caused by anesthesia may cause tonic stretch of this muscle, explaining the tonic activity of the MSAs.

Phase tuning of individual MSAs reflects a combination of stretch sensitivity and top-down modulatory drive. Stretch-related phase tuning is set by single-unit response properties (units with more velocity sensitivity will fire earlier in each lick cycle, compare Figure 3f-h vs. Figure 3d-e) as well as complex stretch fields in the jaw musculature. Cycle-by-cycle modulatory drives may include contraction-linked (i.e. ‘α-γ coactivation’) drives and/or other cyclic modulatory activity. We see three general types of MSA phase tuning (Figure 2e*–*f): (1) units that fire during the opening swing, just before tongue protrusion; (2) units that fire around maximal jaw opening and maximal tongue protrusion; and (3) units that fire after tongue retraction during the closing swing. Our results reveal a relationship between MSA phase tuning, location, and kinematic tuning, as well as jaw muscle activity. For example, ‘jaw open – tongue out’ units (Figure 3i–j; Supplemental Figure 7c), located in *temporalis* or *anterior masseter*, are tuned to maximal jaw opening phases, while ‘jaw open – tongue in’ units (Figure 3k–l; Supplemental Figure 7c), located in *posterior* or *ventral masseter*, are tuned to jaw closing phases. Phase tuning peaks for both groups coincide with peaks in muscle activity (Supplemental Figure 4d). Our results suggest that MSAs may be used to monitor (Figure 6c,f) and/or coordinate progression through the stages of the lick cycle.

### Flexible, task-driven tuning of MSAs

Past work in humans has shown that MSAs are modulated based on higher-order task goals^13,31^, including during the preparation of upcoming movements^32^. We studied MSA activity in the context of motor sequence performance. Licking behavior is controlled by hierarchical somatomotor cortical circuits. Primary sensory (tongue–jaw S1, S1TJ) and motor (tongue–jaw M1, M1TJ) cortices encode sensory stimuli and drive lick movements, respectively^33–35^. Tongue–jaw premotor (anterolateral motor, ALM) cortex is involved with planning and initiating licking as well as online correction of ongoing lick movements^36–38^. During the directed lick sequence task, prior work shows that S1TJ and M1TJ encode cycle-by-cycle orofacial kinematics, while ALM encodes sequence-level kinematic parameters (such as the intended lick angle) as well as higher-order task parameters (such as the sequence identity, sequence progress, and reward state)^3^ (Figure 6g).

We found mixed representations of kinematic and task information among jaw MSAs during lick sequence performance (Figure 6). MSAs reported kinematics within and across lick cycles; kinematics could be decoded from MSA ensembles on both time scales (Figure 6). MSAs were also tuned to progress within the sequence (Figure 4), and their tuning could depend on the reward-context of the lick (Figure 5). These neural dynamics parallel the multiscale dynamics seen in sensorimotor cortex (Figure 6g). Our data suggest that cortical/subcortical regions enact flexible, task-level control over fusimotor neurons and jaw MSAs via direct projections^27^ or through premotor circuits.

Flexible MSA tuning may help the animal achieve kinematic and/or mechanical goals during sequence execution. For example, we found order-tuned units showing greater (Figure 2a, Unit 6; Figure 4b) or weaker (Figure 2a, Unit 4; Supplementary Figure 8d) activity during the early (i.e. first through third) licks of the sequence. We similarly saw units showing greater (Figure 5b) or weaker (Figure 5d) activity during drive compared to drink licks. MSAs and lower motor circuits are embedded in nested closed loops, and these MSA dynamics across the sequences co-occured with changes in the muscle contraction (i.e. α motor neuron) drive (Supplemental Figure 10).

The task-driven feedback dynamics we identified may reflect changing requirements for jaw stiffness/damping^19^, movement robustness, and/or somatosensory information across sequence execution. For example, it is possible that the first sequence lick is more ‘reach-like’ as the animal searches for the port, while drink licks after reward delivery represent more ballistic, rhythmically patterned movements. Task-driven modulation of feedback may, for example, allow the system to change its capacity to reject perturbations while achieving similar movement trajectories. Such changes should be identifiable in both the sensory feedback and motor output signals, as we observed (Supplemental Figure 10h).

Real time tuning of feedback gains in response to evolving operating conditions (“gain scheduling”) is a standard feature of engineered control systems^39^. Circuits in the central nervous system similarly tune feedback gains^40^, including the gains of spinal reflexes directly downstream of MSAs^24,25,41^. Our work suggests that tuning of first-order MSA inputs is a feature of adaptive motor control. Given the long-standing interest in biological motor control solutions for engineering and artificial intelligence research^1^, our results offer insights into how adaptive sensation can be utilized to build complex behaviors.

## Acknowledgements

We thank Duo Xu, Rajan Dasgupta, Mingyuan Dong, Montrell Vass, William Snider, Ki Yoon Nam, Jong Cheol Rah, Bilal Bari, and Yuxi Chen for assistance in data collection; Samuel Sober for Myomatrix arrays. This work was supported by NIH grants 1F32MH120873-01, 1R01NS104834-01,1RF1NS131984-01, R01NS109237, U24NS126936, and the Kavli Foundation.

## Author contributions

W.O., V.C., and J.J.K. performed experiments. W.O. developed analysis code and analyzed data with input from all authors. D.H.O., and N.C. conceptualized and guided the experimental design and data analysis. W.O. and D.H.O. wrote the paper with input from all authors.

## Materials and Correspondence

Requests for materials and correspondence should be directed to D.H.O.

## Methods

### Mice

All procedures were in accordance with protocols approved by the Johns Hopkins University Animal Care and Use Committee (protocols MO018M187, MO21M195, and MO24M185). Mice were housed in a room on a reverse light-dark cycle, with each phase lasting 12 h, and maintained 20–25° C and 30–70% humidity. Before surgery, mice were housed in groups up to five, but afterwards were housed individually. All mice used in this study were obtained by mating Cre lines with wildtype (Jackson Labs 000664; C57BL/6J) or Ai32 (Jackson Labs 012569; B6;129S-Gt(ROSA)26Sor^tm32(CAG-COP4*H134R/EYFP)Hze^/J) lines. Mice used in this study included two Advillin-Cre (Jackson Labs 032536; B6.129P2-Avil^tm2(cre)Fawa^/J) females, two Calb1-Cre; Pvalb-Flp (Jackson Labs 028532; B6;129S-Calb1^tm2.1(cre)Hze^/J, Jackson Labs 022730; B6.Cg-Pvalb^tm4.1(flpo)Hze^/J) females, four Dbh-Cre (Jackson Labs 033951; B6.Cg-Dbh^tm3.2(cre)Pjen^/J) mice (two males and two females), one TH-Cre (Jackson Labs 008601; B6.Cg-7630403G23Rik^Tg(Th-cre)1Tmd^/J) male, and two wildtype mice (one male and one female). Mice were 2–8 months old at the start of behavior training, and training and testing sessions typically lasted 2–3 months.

### Surgery

Mice underwent surgery for the implantation of a headpost and microdrive. For surgery, mice were anesthetized with 1–2% isoflurane and held on a heating blanket (Harvard Apparatus). Ketoprofen was injected i.p. to reduce inflammation, and lidocaine or bupivacaine as a local analgesic was injected under the scalp at the start of surgery. The skin and periosteum above the skull were removed. A small circular craniotomy was made with a dental drill over the anterior cortex for the implantation of a grounding pin. The ground pin was either a small stainless steel screw or a gold male pin soldered to a short length of silver wire fixed to the skull with dental cement. A thin layer of metabond (C & B Metabond) was applied to cover the surface of the skull to fix the headpost onto the skull. The custom-made metal or 3D printed headposts were positioned over the lambda suture and feature a recessed opening, allowing access to the skull posterior to the lambda suture.

A ∼2 mm diameter craniotomy was made centered at −1mm caudal and +1.1 mm lateral to lambda, near the border of the cerebellum and the inferior colliculus. All implantations occurred on the animal’s left side, to target MSA neurons innervating the left jaw musculature. To map the location of MEV within this craniotomy, tungsten recordings were performed to search for jaw movement related LFPs. The craniotomy was covered with sterile PBS, the dura was carefully removed using a tungsten needle, and a tungsten recording electrode (0.5 MΩ, WPI) was lowered into the brain using a micromanipulator (Sutter instruments). The differential signal between the recording electrode and a bath reference (AgCl pellet) was amplified (DAM80, WPI) and monitored on an oscilloscope (Tektronix) and through an audio monitor (A-M Systems). Characteristic LFP responses to ∼1 Hz manual jaw movement were typically found ∼-2.5–4 mm below the surface of the brain.

Custom-built microdrives containing eight tetrodes (nichrome wire, 100–200 kΩ, Sandvik) and an optical fiber (0.39 NA, 200 μm core) were implanted at the identified location. The microdrive was slowly lowered to ∼0.5–1 mm above the identified dorsoventral position, then fixed to the skull with dental cement. The recording was grounded to the grounding pin before shielding the microdrive with plastic/aluminum foil housing. Mice were given a minimum of 1 week of recovery before water deprivation.

Three mice received virus injection (200–750 nL, ∼5 nL/s injection speed) into the MEV or the nearby locus coeruleus. Viruses (Addgene) included AAV.EF1a.DIO.hChR2(H134R).eYFP, AAV.hSyn.Con/Fon.hChR2(H134R).EYFP, or AAV.mCherry.ChR2.

### Sequence task behavior

The behavior apparatus, task, and training have been previously described^3^. Briefly, the task was controlled by an Arduino-based system and custom MATLAB software. This system moved a two-axis motorized (LSM050B-T4 and LSM025B-T4, Zaber Technologies) lick port, monitored lick port contact registered via a conductive lick detector (Svoboda lab, HHMI Janelia Research Campus), delivered trial start auditory cues (a 0.1s long, 65 dB SPL and 15 kHz pure tone) and a constant masking auditory stimulus (white noise and previously recorded motor noise), delivered water rewards (∼2–3 μL) via the lick port, implemented user control of task parameters, and logged behavior control and performance.

During the task, the port moved through an arc of seven locations specified via a polar coordinate system. The origin of this system was referenced to the mouse’s incisors, so that these positions were symmetrical to the mouse’s midline with equal spacing in arc length. The radius and total arc distance of the positions were adjusted to control task difficulty, with smaller radii/lengths used for early training. Difficulty was gradually increased over 7–15 training sessions to reach performance of ∼100–400 trials in ∼1 hour behavior testing sessions. Mice performed the task in the dark without visual cues about the port position. Ventral whiskers (macro and microvibrissae) were regularly trimmed to prevent any whisker-based localization of the lick port.

### Behavior testing, unit recording, and unit characterization

Prior to behavioral training, the microdrive was slowly advanced while the presence of jaw movement-related LFPs was assessed to place the chronic recording in MEV. The mouse was water-deprived, and then received behavior task training. Toward the end of training, the mouse’s left cheek was dehaired using Nair depilatory cream, and the ventral whiskers were trimmed. After 1–2 further training sessions, mice underwent behavior testing and recording.

For a given recording session, high-speed video and tetrode electrical activity were recorded while the mouse performed the sequence task for ∼0.75–1 hr. Extracellular voltages were amplified and digitized at 30 kHz via an RHD2164 amplifier board and acquired via an RHD20000 system (Intan Technologies) without filtering. The microdrive was typically advanced ∼100um per day across a set of testing sessions.

After behavior testing, the animal was induced under light isoflurane anesthesia (∼0.5–1%) for a panel of passive characterization stimuli. The depth of anesthesia was adjusted to maintain a breathing rate of ∼2 Hz, and the passive stimulus panel typically lasted ∼0.75 hr. The jaw was first passively moved with a wooden stick. ∼1 Hz movement was applied in multiple directions (up-and-down and side-to-side in both directions). In some sessions, faster ‘lick-like’ movements were applied to the jaw. Extracellular recording and high-speed video were acquired during movement. Movement was timed to a 1 Hz auditory tone delivered by the Arduino. The Arduino delivered cues to the neural recording software synced to this tone, allowing movement to be registered to the neural and video recording.

To record unit cheek probing responses, a camera (Thorlabs CS165MU1, fitted with an 8 mm fixed focal length lens, Thorlabs MVL8M23) was positioned above the mouse to view the left cheek via a silver-coated mirror. Synchronous triggers, synced to an auditory tone, were sent to the recording system and the camera, which captured 3 frames (1 ms exposure, 10 frames per second). Cheek probe stimulation was manually timed to the tone. Inter-stimulus intervals were either 5 s or 2 s. Stimuli included a blunt wooden stick (∼2 mm diameter), a blunt metal piece sheathed in rubber (∼1.6 mm diameter), and von Frey hairs (0.6 g - 2.0 g). Before acquiring data, the cheek was searched during audio monitoring of extracellular responses to localize unit responses. During mapping acquisition, the cheek was searched with multiple passes over the entire cheek within a session. During most sessions, high-speed video was acquired during the probing to localize the stimulus on the cheek and exclude stimuli that caused jaw movement.

### Videography, tracking, and kinematic analysis

As previously described^3^, high-speed (400 Hz, 0.6 ms exposure time, 32 µm per pixel, 800 × 320 pixels) dual-view video was acquired via Streampix 7 software (Norpix) and a single camera (PhotonFocus DR1-D1312-200-G2-8 camera) fixed with a x0.25 telecentric lens (55–349, Edmund Optics). Side and bottom views were simultaneously acquired via a silver-coated mirror fixed under the mouth region. The mouth was illuminated via an 850 nm LED (LED850-66-60, Roithner Laser) passed through a condenser lens.

3D jaw and tongue keypoints were tracked from the dual view video using DeepLabCut^42^. A network was trained to detect three points on the chin (forming a small triangle on the distal chin) and two points on the tongue (the base and tip of the visible tongue) from both views. Training data included 1070 manually labeled frames from 9 mice. The frames were selected from videos of mice performing the sequence licking task and mice receiving manual jaw movement under anesthesia.

We then mapped kinematic tracking data from the various animals into a common reference space. Resting frames were identified as those in which movement velocity did not exceed 3.2 mm/s for any jaw keypoint. A rigid transformation (rotation + translation) was calculated mapping the session average jaw rest position onto a common reference position, and this transformation was applied to all tracked keypoints.

The mean (centroid) of the three jaw-tracked keypoints was taken as the jaw position. Tongue length was taken as the 3D Euclidean distance between the visible tongue base and tip keypoints. The angle (in radians) of the base-tip vector from the vertical in the XY plane was taken as tongue θ, and the angle of the base-tip vector from the horizontal in the YZ plane was taken as tongue φ. Tracking data were smoothed with a 3rd-order Savitzky-Golay filter with a window size of 27.5 ms.

Jaw setpoint *x* was extracted by segregating the data into single lick events (using phase decomposition, see below), estimating setpoint *x* at phase=0 rad. (maximal jaw opening) as the lick median jaw *x* value, and linearly interpolating for the other time points^43^.

Jaw–tongue coordinate kinematic axes were identified using Singular Value Decomposition (SVD) of jaw–tongue features extracted from the keypoints in each video frame (time bin size = 2.5 ms). For each session independently, data were first limited to licks passing a jaw *z* range of 0.5 mm (see Phase decomposition) and z-scored. SVD was performed (scipy.linalg.svd). The right singular vectors of the SVD were taken as the coordinate axes, the left singular vectors were taken as the weights onto these axes, and the singular values represented the relative variance explained by the axes.

Kinematic features were extracted from the video data by SVD using Facemap^17,18^. We used a spatial binning parameter of 10 pixels. Weights for 500 features calculated via SVD of the raw frames and for 500 features calculated via SVD of the motion energy between frames were concatenated. Weights for all 1000 features were used for the prediction of neural activity (see below).

### Electromyography

Temporalis and masseter muscle bipolar EMG recordings were taken using either custom-made fine-wire hook electrodes (PFA-coated tungsten wire, A-M Systems 795500) (n = 4 sessions from 2 animals) or high-density Myomatrix EMG arrays^44^ (n = 2 sessions from 2 animals). Recordings were grounded to implanted pins in the skull, digitized at 30 kHz via a 16 channel bipolar input headstage (RHD2216, Intan Technologies) and acquired via an RHD2000 system.

### Histology

After the conclusion of testing sessions, electrolytic lesions were made under anesthesia in one tetrode channel. Mice were transcardially perfused with PBS, followed by fresh 4% PFA. The brain was dissected and post-fixed in 4% PFA overnight. The midbrain was sectioned on a vibratome (50–100 μm). Sections were then immunostained for Parvalbumin: sections were permeabilized in PBS + 0.5% TritonX-100 (PBT), incubated in 5% donkey serum + 1:1000 rabbit anti-parvalbumin (Abcam ab11427) + PBT (overnight at 4°C), washed in PBT, incubated in 1:1000 anti-rabbit IgG-488 or 594, washing in PBT, and mounted in Aqua-Poly/Mount (Polysciences). MEV was identified via location and characteristic PV+ large diameter cell bodies, and the lesion location was confirmed (Supplementary Figure 1b).

### General data analysis and plotting

Analysis was performed using MATLAB (2023a), Python 3.10, and ImageJ. Large Language Models (LLMs; ChatGPT and GitHub Copilot) were used to suggest some analysis code, but all code was carefully reviewed by the authors. Boxplots show the median as a center line, quartiles 1 and 3 as box limits, and ±1.5x the interquartile range as whiskers (outliers are not shown). Error bars are defined in figure legends.

### Spike sorting and unit inclusion criteria

We used Kilosort2 for initial spike sorting. Extracellular recording data were high-pass filtered at 500 Hz and across-channel-median subtracted (common average referencing) before passing into Kilosort. Spike sorting results were manually curated in Phy and MClust. We used three quality control metrics to select single units (Supplementary Figure 1c–e): (1) interspike interval violation rate (ISI VR), (2) false positive rate (FPR), and (3) signal-to-noise ratio (SNR). We saw high-frequency bursts of MEV unit activity with inter-spike intervals of ∼1 ms (Supplementary Figure 1e). We chose a 1 ms refractory period size and excluded units with ISI VR > 2%. FPR was estimated using a published method^45^, and we excluded units with FPR > 15%. SNR was calculated as the unit waveform amplitude divided by the channel standard deviation for the channel with the largest waveform amplitude. Units with SNR < 4 were excluded.

In anesthetized animals, the presence of robust jaw movement-related multi-unit activity is a hallmark of MEV in the local region (Supplementary Figure 1a). For a given session, to select recordings from tetrodes located in MEV, we calculated the multi-unit autocorrelogram (ACG, +/−5 s with 10 ms bin size) of each recording channel during 1 Hz manual jaw movement. Multi-unit spikes were detected using a threshold-crossing method, with the threshold set at 5.93 times the median absolute deviation of the channel. Channel recordings were classified as within MEV if a peak detection method (scipy.signal.find_peaks, rel_height threshold of 0.5, prominence threshold of 0.5) identified peaks in the ACG at both 1 and 2 s lags (+/− 0.2s). Tetrodes with any channel passing this criterion were classified as MEV recordings for the session.

MEV units were further tested for movement-related modulation in the awake animal using the ZETA test^46^ (Supplementary Figure 1f–h). This test references spike times against a behavioral event, collects cumulative referenced spike times within a test window, and compares the cumulative spike times array against a null distribution in which the event times are shuffled against the spike times. We referenced unit spike times against the time of lick contact at the central port, using a 1 s test window. Units with a significant (p<0.05) ZETA (Zenith of Event-based Time-locked Anomalies) were classified as movement-modulated. Most within-MEV unit recordings are modulated by active movement (103/116 unique units, Supplementary Figure 1h). Units passing the ZETA criterion sometimes appeared to show little task modulation when spikes were aligned in time, yet subtle movement-related activity was revealed when spikes were aligned on lick cycle phase (Supplementary Figure 1g).

Across sessions, all single units found on the same tetrode were compared. Units with similar single-unit ACGs, waveforms, and activity during the sequence licking task were classified as putative duplicate units (seen across 2–4 sessions). For analyses of the single-unit dataset, only one session was kept for each putative duplicate recording. If a cheek response field was found for one session, that session was kept. Otherwise, the session with the highest SNR was chosen. When cheek response fields were found in multiple sessions, the response fields were consistent across sessions, and the highest SNR session was kept. Analyses using ensemble models (Figure 6) compare information across small populations of units rather than across individual units. Duplicates (representing a small fraction of the overall population) were not removed for ensemble analyses.

Lastly, prior reports indicate a subset of MEV neurons are low-threshold mechanoreceptors (LTMRS) innervating the teeth or whisker pad^47–49^. Units with tooth-region probe response fields showed low firing rates during the task (Supplementary Figure 3). Therefore, MEV units with low mean firing rates (<22 Hz, 26/104 unique units) calculated across the entire awake recording were considered potential low-threshold mechanoreceptors (LTMRs) and were excluded from the analysis.

### EMG signal processing

EMG recording signals were bandpass (400Hz - 3kHz) filtered with a 2nd-order Butterworth filter. Electrical lick port contact artifacts were removed by censoring data in 5 ms windows centered on contact times and linearly interpolating over the gaps. Signals were then rectified and smoothed with a 10 ms standard deviation Gaussian kernel.

### Cheek probe response maps

Probe locations were manually identified in ImageJ. Image stacks for each session were registered to a common cheek image (Supplementary Figure 2c,j) using manually selected keypoints (fitgeotrans in MATLAB, projective transformation). 0.25 s test windows were centered on the time of auditory tones. Single-unit spikes were binned (10 ms bin size), spike rates (spikes/s) were smoothed with a 20 ms std Gaussian kernel, and the maximum spike rate was taken as the unit test window response. Responses were combined across stimulus types (i.e. blunt probes and von Frey hairs). For sessions with simultaneous jaw video recording, responses from test windows with >0.32 mm movement were excluded.

### Phase decomposition

Lick-cycle phase was decomposed from the jaw kinematics via methods previously used to decompose whisk cycle phase from whisker kinematics^43,50^. Jaw *z* data were first smoothed with a 50 ms window moving median filter and then detrended with a Butterworth bandpass filter (2–20 Hz). Lick cycle phase was taken as the angle of the Hilbert transform of this signal. Rare short stretches in which the decomposed phase ‘doubled back’ (i.e. quickly decreased and increased) were identified and replaced with linearly interpolated values.

Sequence progress was identified by aligning sequence task data on the phase value of the start of the first lick with a port contact, unwrapping phase assignment across the trial, and dividing by the radian value of the time of water delivery. This assigns an arbitrary value of 0 to the start of the first contact lick and 1 to the time of water delivery. Phase assignments are unreliable for the stationary jaw and for low amplitude licks. Cycles in which the jaw *z* range did not pass a threshold of 0.5 mm were excluded from the analysis.

### Lick and trial selection

For lick cycle analyses (kinematics, EMG, unit responses), licks within a session were ranked by the range of jaw *z* movement, and the top 25% of all session licks were kept (636–2174 licks per session).

For sequence trial analyses, a custom trial selection algorithm was used to choose the best matched trials across animals and sessions. For each trial, jaw *z* and *x* data were interpolated using unwrapped sequence phase (in rad.) onto a common array (the first through seventh licks, with 20 points per lick cycle) and concatenated into a single array. Within a sequence type (i.e. left-to-right), the median across-animal, across-session sequence array was found. Within a session, 50 left-to-right and 50 right-to-left trials with the smallest Euclidean distance from this median array were selected.

### Averaging and bootstrapping

Kinematics and EMG signals were averaged in phase (scipy.stats.binned_statistic) with a bin size of pi/20 radians. Unit phase histograms were calculated by counting spikes in each pi/20 radian bin and dividing by the total real time spent in each phase bin. Phase-aligned trial histograms were calculated similarly, binning spikes on unwrapped phase assignments across the trial (for −1.5 to 9.5 cycles from the start of the first contact lick). Error estimates (standard deviation and 95% confidence intervals) were calculated via bootstrapping (1000 resamples). For within-session estimates, resampling with replacement was done on the trials. For across-animal, across-session averages, we employed hierarchical bootstrapping; on each pass, resampling with replacement was done at the level of the animal, then session, then trial (e.g. Figure 2a, *z* and *x*), or animal, then session, then lick (e.g. figure 2d).

For 2D tuning surfaces and associated 1D tuning curves, we calculated 30 non-uniform bins for each parameter to more evenly distribute observations across the bins, as previously described^50^. The sigmoid function parameter k was selected as the inverse of 0.3-0.4 times the maximum absolute value of the variable for each curve in order to ensure adequate coverage of the relevant kinematic space. For neuronal tuning surfaces, binned spike counts (scipy.stats.binned_statistic_2d) were divided by the total real time spent in each bin. For lick contact probability surfaces, the number of bin observations occurring with lick port contact was divided by the total number of bin observations. Surfaces were smoothed with a Gaussian 2D filter (kernel std. width of 1.5 bins). Bins with fewer than 10 observations were excluded (colored white on 2D surfaces).

### Phase tuning analysis

We segregated units based on the strength of phase tuning. Some units showed bimodal phase tuning responses, which posed a challenge for tests that use vector strength (i.e. the Rayleigh test). We therefore identified units in which the spike counts in 2.5 ms bins had significant mutual information (MI) with lick cycle phase during task performance. Data were limited to task performance windows (0–16 lick cycles from the start of the first contact lick on each trial) because phase assignments are inaccurate when the jaw is not moving. The spike count distribution was *P*(*X* = *x*), *x* ɛ {0, 1, 2, … *n*}, where *n* is the unit maximum spike count (*n* ≤ 5). The phase distribution *P*(*Y*) was estimated by binning phase (*-π* to *π* rad.) into 20 uniform bins. The joint distribution *P*(*X* = *x*, *Y* = *y*) was estimated similarly, and the MI^15^ was computed as:

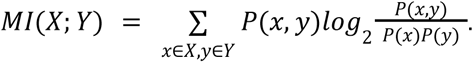

We calculated 90% confidence intervals for MI under the null hypothesis of no correlation by shuffling the spike counts against the phase values for 1000 iterations. We further calculated normalized MI by dividing MI by the spike count entropy:

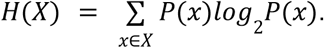

Units for which MI was above the 90% CI under the null hypothesis and for which normalized MI > 0.01 were deemed significantly coupled to phase.

Phase tuning peaks were identified for phase-coupled units and for EMG recordings. Unit phase histograms (or phase-averaged EMG signals) were concatenated on both ends with repeated copies (for smoothing over the ends), smoothed (3^rd^ order Savitzky-Golay filter with a window size of 27.5 ms), and normalized by the maximum value. A peak detection algorithm (scipy.signal.find_peaks with a prominence threshold of 0.1) was run, and the phase value of the peaks were kept. Phase-coupled units and EMG recordings showed either one or two peaks. These were ordered as primary and secondary peaks based on peak height. To calculate the probability of tongue protrusion out of the mouth as a function of lick cycle phase, a binary vector representing bins with a tongue length > 0.5 was averaged based on phase across all animals and sessions.

### Encoding and decoding linear regression

Encoding and decoding linear regression models were fitted on binned (2.5 ms bin size) kinematic and spike rate data. Unit spike counts were binned, and spike rates (spikes/s) were smoothed with a Gaussian kernel. The kernel width (std = 25 ms) was chosen to maximize overall encoder model performance (Supplementary Figure 4b). Unit-coordinate axis correlations (absolute value of the Pearson correlation *r*) were calculated between axis projection weights and smoothed spike rates. The 3 best correlated axes were used for linear models.

Linear models were fitted using sklearn. Unless otherwise noted, model predictive performance was calculated using Monte-Carlo cross-validation (MCCV) on the trials. On each pass (resample number = 1000), 10% of trials were randomly chosen as the testing set, models were fitted on the training set (remaining 90% of trials), and performance was evaluated on the testing set. Reported cross-validated *R*^2^ or Pearson’s r values are the mean of the resample model performances. Pearson’s r is used for phase decoder performance (Figure 6f) since *R*^2^ is a poor goodness-of-fit measure for circular data.

For significance testing of the difference between two models (Figure 3n; Figure 5f), performance was compared within each resample. For example, in Figure 3n, on each resample, jaw+tongue encoder performance was subtracted from jaw encoder performance. If the 95%CI of this comparison was greater than 0 (equivalent model performance), the performances were significantly different. For Figure 5f, if the 95% CI did not include zero, the performances were significantly different.

For comparison of encoder model performance between drive and drink conditions (Figure 5), the drive licks explored a wider movement space than the drink licks. Therefore, we limited drive lick data to samples within the span of the drink lick jaw movement space. For each session, 100 uniform bins were found for the range of jaw *z* and *x*. We then identified *z*, *x* 2D bins with > 10 observations during drink licks, and excluded samples in the drive lick data that did not match these bins. SVD was performed as above on the combined (drive and drink lick) restricted data and used for fitting encoding models.

Ensemble decoding models were fitted to linearly predict single kinematic or task parameters using smoothed (25 ms std. Gaussian kernel) spike rates from simultaneously recorded units. Model predictive performance was calculated using 5-fold cross-validation. Nested ensemble decoders missing subsets of these units were analyzed using an iterative unit dropping strategy: for each round, each unit was dropped in turn, and the 5-fold c.v. predictive performance was found for each nested ensemble. The nested ensemble with the largest performance loss (i.e. the ensemble lacking the top-performer unit) was used for the subsequent round.

### Boosted trees models

Boosted trees encoding models using kinematic regressors to predict spike rates in 2.5 ms bins were fitted using XGBoost with a mean squared error objective. Hyperparameters were chosen to maximize cross-validated performance; we used 25 gradient-boosted trees and a 0.2 learning rate *eta*. We used a subsampling ratio of 0.2. Predictive performance was calculated using 10-fold cross-validation on the trials.

### Binary decoder analysis

Binary decoders were fitted to classify lick position or order for licks to two selected port positions (L3 and R3, or L1 and R1). Within each port position, we sub-selected licks to match kinematics using a custom algorithm. An array containing the mean jaw *z* and *y* positions was found for each lick, and the pairwise Euclidean distances between all lick arrays were calculated. We then iteratively chose the best-matched pair, removed this pair, and chose another pair. 10 pairs were chosen for each position, yielding a total set of 40 licks.

For each unit, we assessed the ability of unit within-lick mean spike count to decode position (e.g. L3 vs. R3) or order (e.g. first vs. last) using the Area Under the Curve (AUC) of a Receiver Operating Characteristic analysis in sklearn (Supplementary Figure 8b,e,h). This analysis varies a test threshold and compares the True and False Positive Rate. An AUC of 0.5 indicates no better than random discriminability, whereas an AUC of 1 indicates perfect discriminability of the condition test distributions. 95% confidence intervals for the AUC scores were found using bootstrapping (1000 resamples) on the trials. If the 95%CI did not include 0.5, decoder performance was classified as significant. The same analysis was performed using mean jaw x, z, and y values (Supplementary Figure 8j). One session in which jaw kinematics could significantly decode lick order was excluded, indicating insufficient performance of the lick selection algorithm. Binary decoders that used EMG activity (Supplemental Figure 10f) were fitted similarly, excepting using within-lick mean rectified and smoothed EMG signals instead spike counts.

## Data availability

All data will be made publicly available by the time of publication, and can currently be provided upon request to.

## Code availability

All code will be made publicly available by the time of publication, and can currently be provided upon request to.

---

**Supplemental Figure 1.**
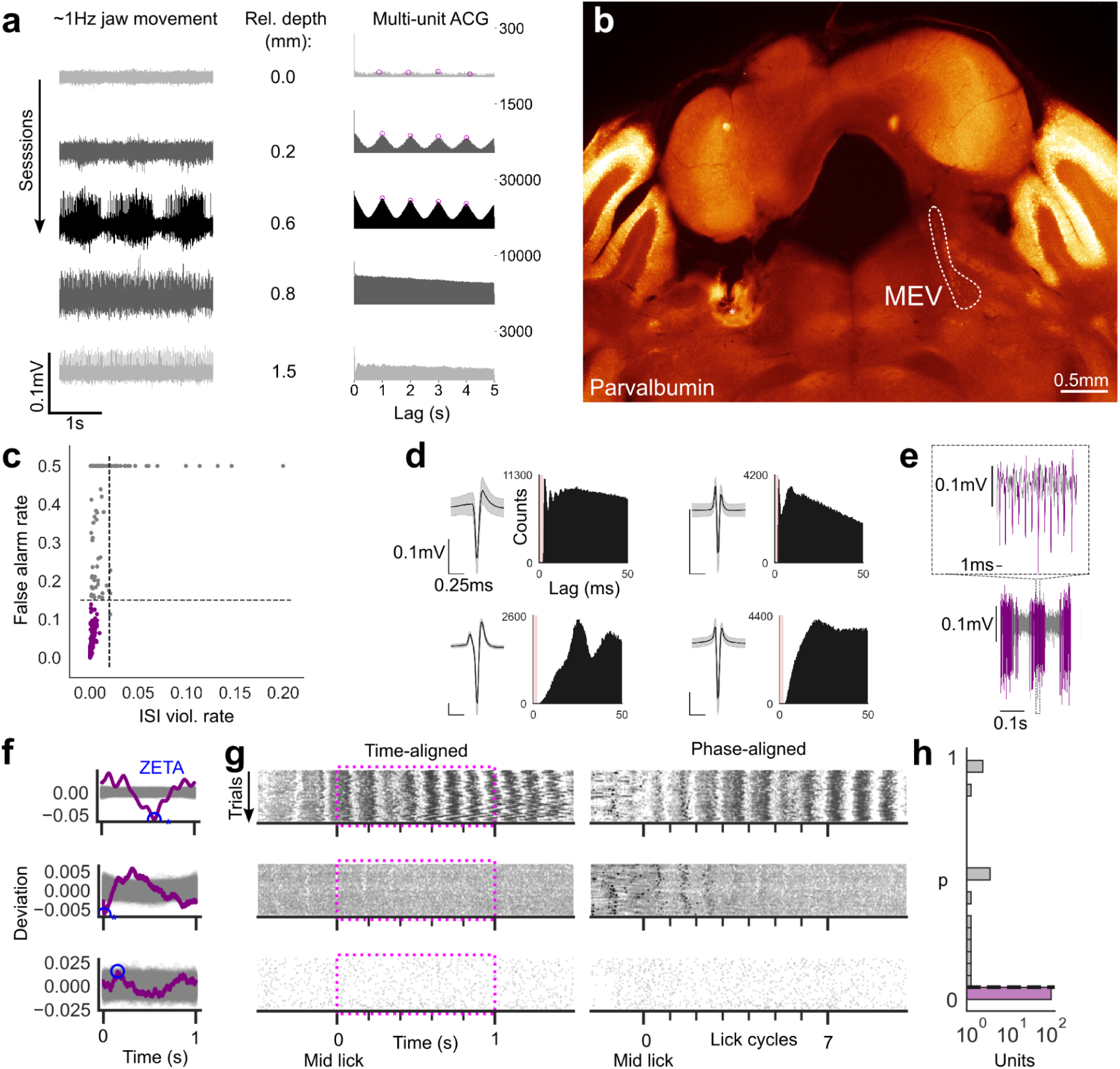
Isolation of jaw MSAs in mesencephalic trigeminal nucleus (MEV) recordings. **a,** Jaw movement-related multi-unit activity under anesthesia is a hallmark of MEV. Left, highpass filtered extracellular recordings during ∼1 Hz jaw movement from a single tetrode channel on subsequent sessions. Middle, microdrive advancement between sessions. Right, tetrode recordings were classified to be in MEV if multi-unit spike autocorrelograms (ACGs) showed peaks at both 1 and 2 s lag. Magenta circles indicate detected ACG peaks. **b,** At the end of a testing session series, electrolytic lesions were performed on a test channel for post-hoc confirmation of recording location. Midbrain section with anti-parvalbumin antibody staining. MEV contralateral to the recording site is indicated. **c,** Single units were kept with interspike interval (ISI) violation rate < 2% and a false alarm rate <15% (Methods). **d,** Representative unit spike waveforms (mean ± std.) and ACGs. Pink box indicates 1 ms refractory period. **e,** Sample extracellular recording with spike detection (purple) for a single MSA. Top panel is an expanded view of the boxed region in the bottom panel. MSAs could fire bursts at rates close to 1 kHz. **f,** ZETA test^46^ for movement modulation during the sequence task for three sample units (Methods), using the lick contact to the center port as a reference event. Purple lines are unit deviation traces, blue circle indicates the maximum deviation (ZETA), gray lines indicate deviation traces under the null hypothesis, blue asterisk indicates significance. **g,** Sequence task trial spike rasters for chosen units aligned either in real time or on lick cycle phase. The ZETA test, performed on real spike times (0*–*1 s window, magenta boxes), could detect subtle movement modulation (compare phase-aligned and time-aligned data for the central row) while excluding non-modulated units (bottom row). **h,** Histogram of p-values for single MEV units (purple, p<0.05). Most (103/116) MEV units were modulated by task movement (x-axis is log-scaled).

**Supplemental Figure 2.**
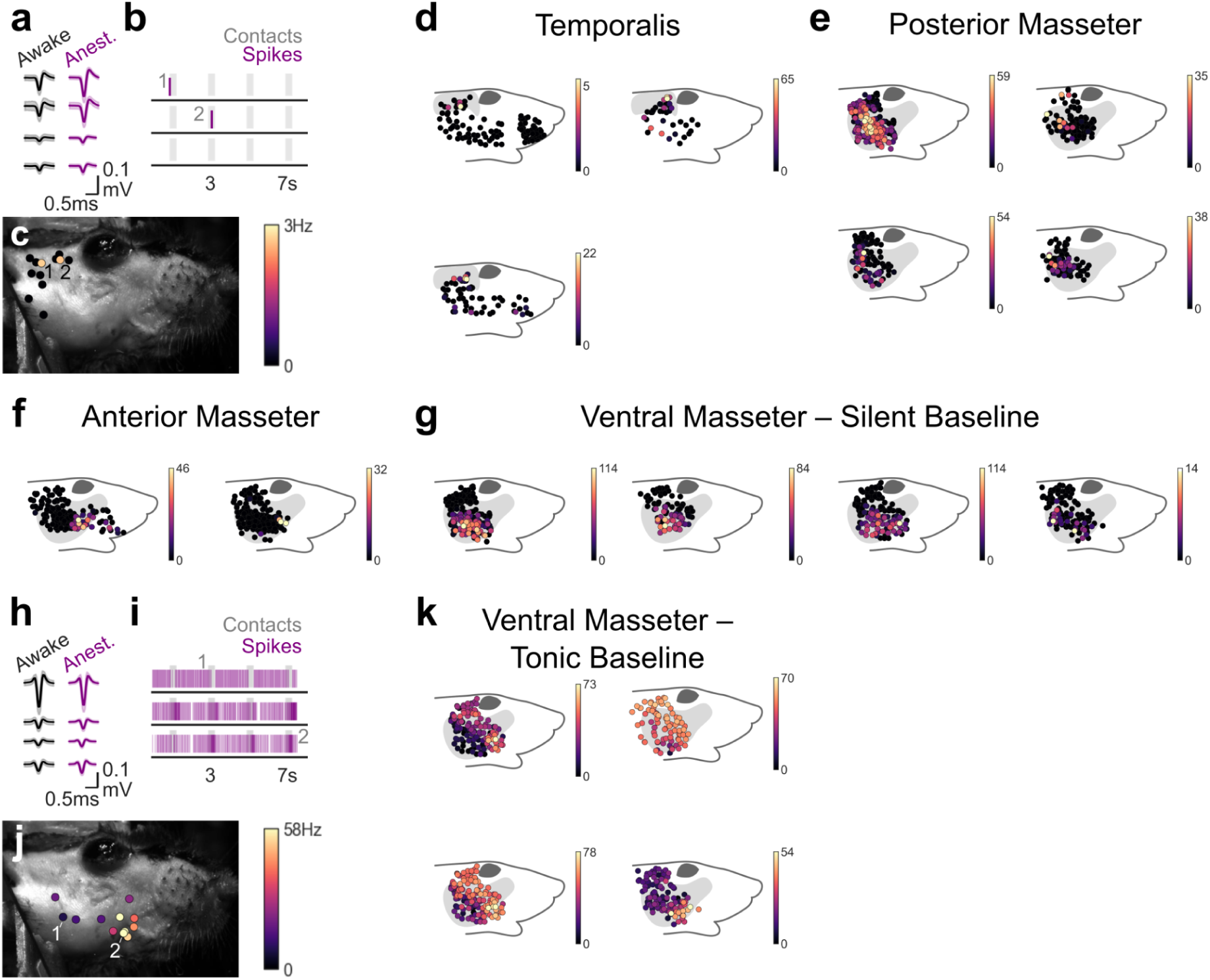
>Localization of jaw MSA muscle of innervation. **a,** Representative unit waveforms (mean ± std.) in awake and anesthetized conditions. **b,** Spike rasters during gentle probing to 12 locations on the cheek. **c,** Maximum spike rates in 0.25 s windows centered on probe stimuli to the 12 locations in b. Locations labeled 1,2 match contact windows labeled in b. **d,** Cheek response maps for 4 units localized to *temporalis* muscle. Color indicates maximum spike rate during the test window. Upper left is the unit shown in a*–*c. **e*–*g,** Maps for four units localized to *posterior masseter*, two units localized to *anterior masseter*, and four units that showed responses localized to ventral masseter in the absence of tonic baseline activity (*ventral masseter – silent baseline*). **h*–*j,** Waveforms, sample contact rasters, and sample cheek map responses for a unit showing tonic baseline activity. Units of this response type fired tonically under anesthesia. These units showed silencing responses to probing in the posterior ventral masseter, and activating responses to probing in the anterior ventral masseter. **k,** Maps for four units with ‘*ventral masseter – tonic baseline*’ responses fields. Upper left is the unit shown in h*–*j.

**Supplemental Figure 3.**
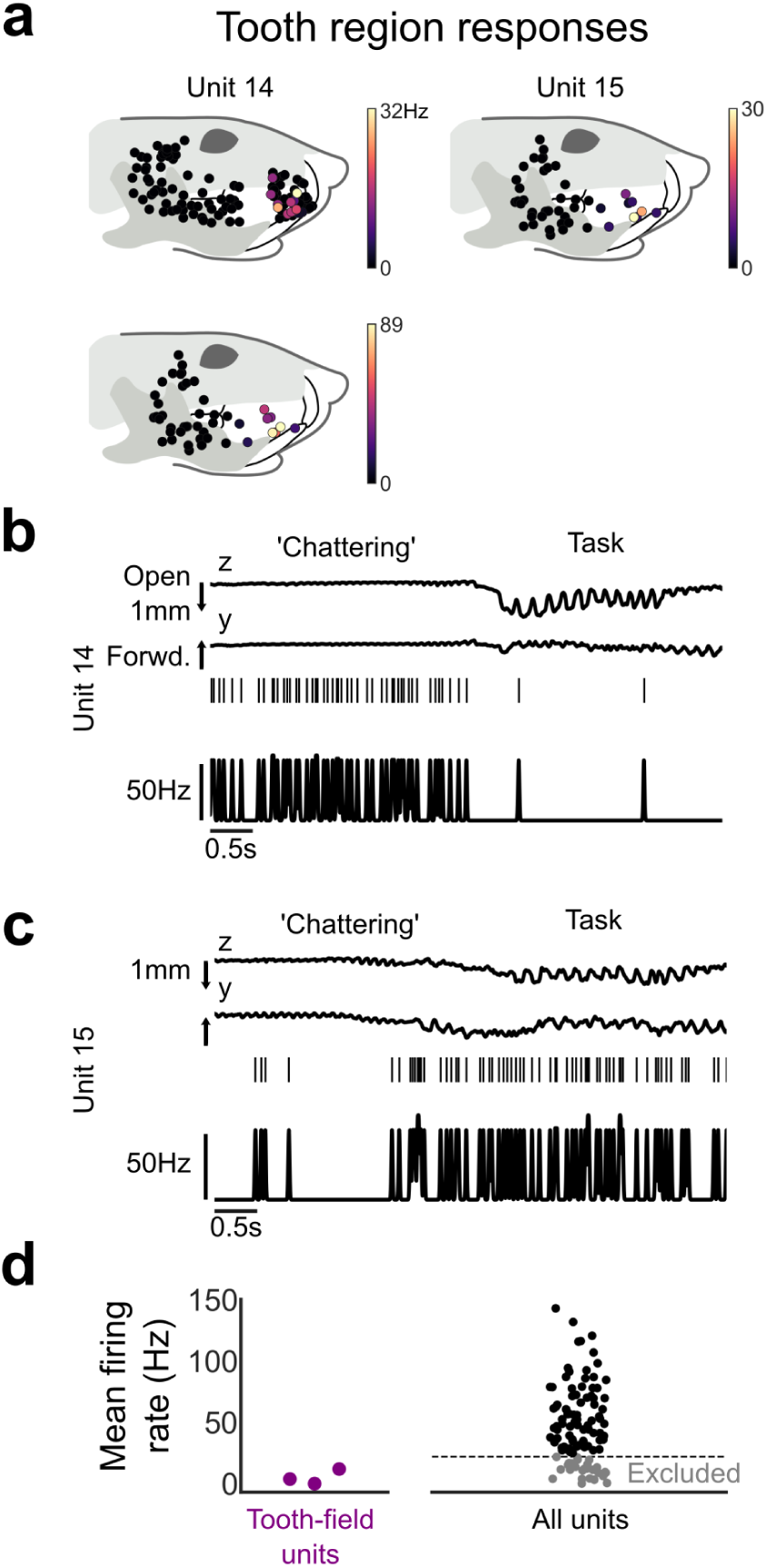
Identification of possible low-threshold mechanoreceptor (LTMR) units in MEV recordings. **a,** 3 units with probe response fields in the tooth/whisker pad region. **b,** Sample unit 14 time-series data (jaw kinematics, spike rasters, and smoothed (7.5 ms std. Gaussian kernel) spike rate before and during task performance. The unit was activated by high-frequency jaw/tooth ‘chattering’ prior to the task, but was not strongly driven by task performance. **c,** Unit 15 showed weaker activation during ‘chattering’, and was moderately activated during the task. Task activity was weakly correlated to jaw movement. **d,** Identified tooth-field units showed overall low mean firing rates during awake behavior performance. We conservatively excluded units with low firing rates from our analysis of jaw MSAs.

**Supplemental Figure 4.**
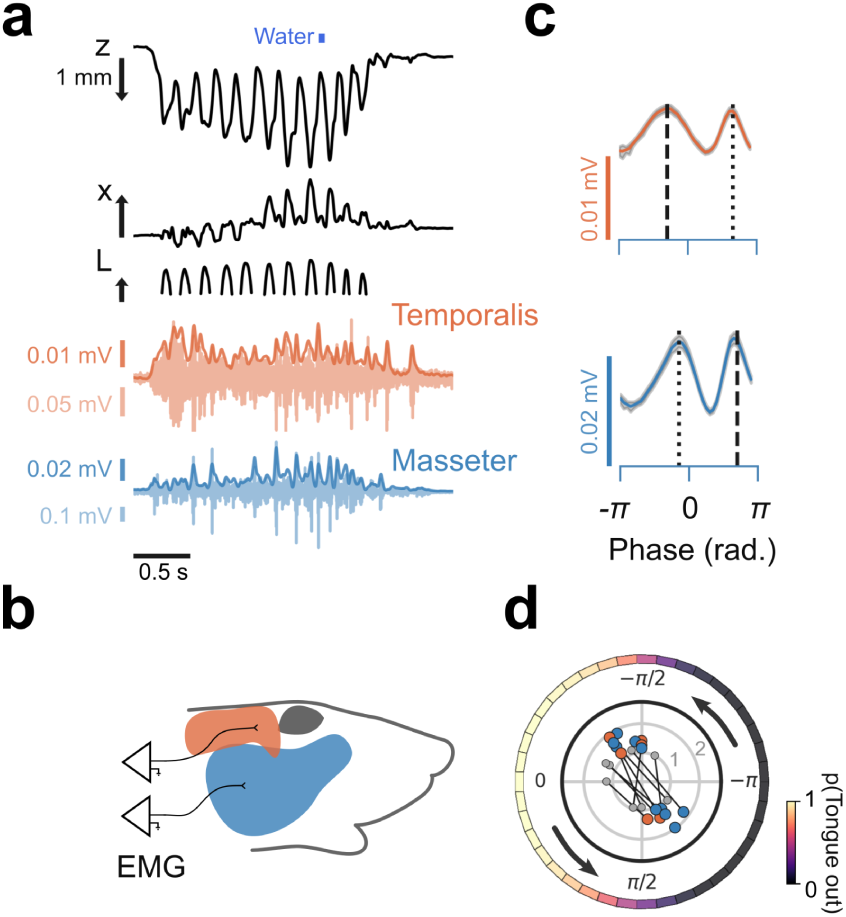
Jaw muscle activity during lick events is associated with tongue protrusions and jaw closing. **a-b,** Electromyography (EMG) recordings from temporalis (orange) and masseter (blue) muscles. Top in a, representative jaw and tongue kinematics. Bottom two rows in a, bandpass filtered EMG (light) and rectified and smoothed (dark) EMG (rsEMG) data. **c,** rsEMG averaged by cycle phase for recordings in a. Dashed line, primary phase tuning peak. Dotted line, secondary peak. **d,** EMG lick cycle phase tuning. Outer, tongue out probability (Methods). Inner, phase tuning peaks (radial axis shows mean-normalized peak height, large circles are primary peaks, small circles are secondary peaks, matching peaks connected).

**Supplemental Figure 5.**
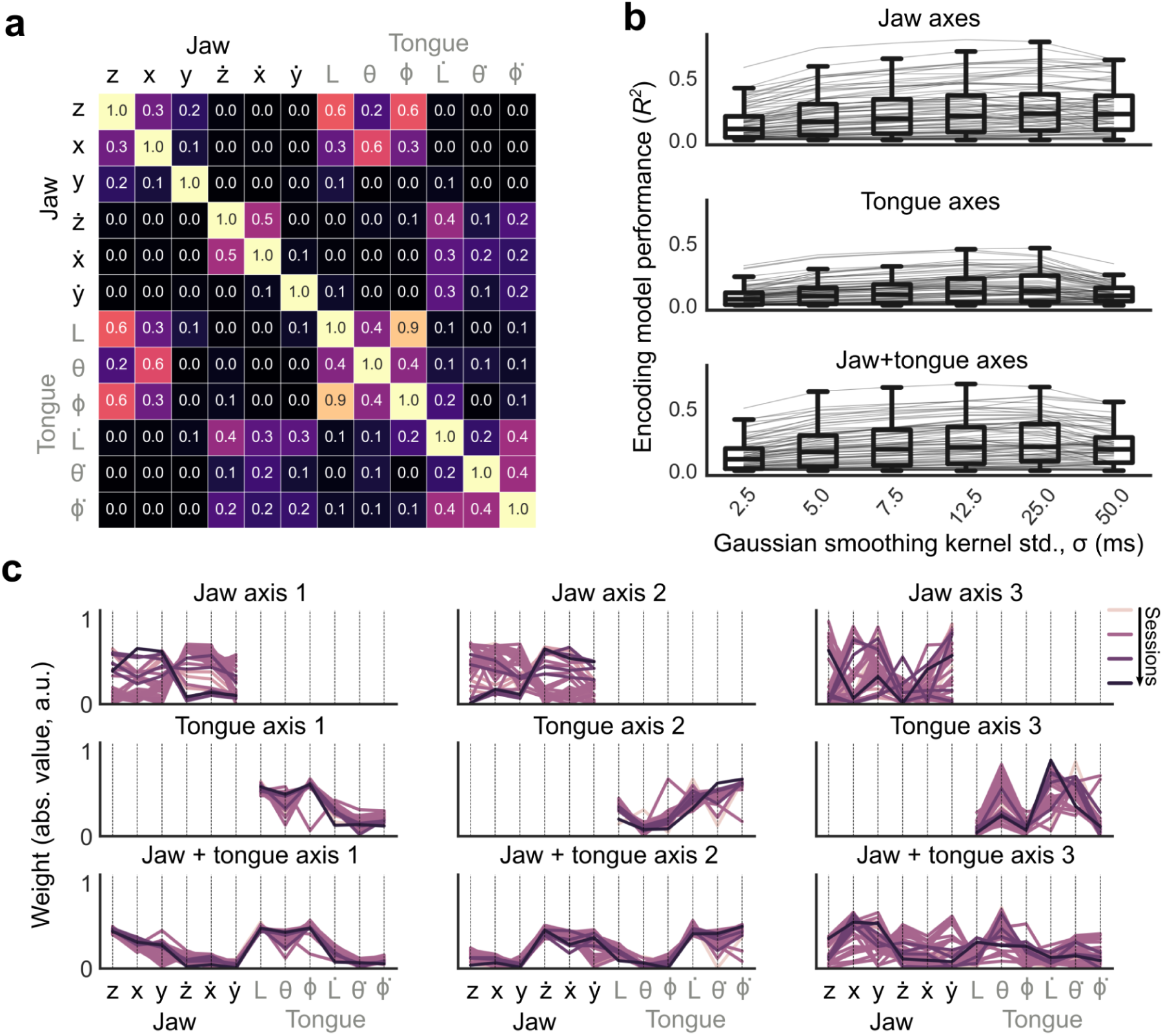
Coordinate axes captured complex orofacial kinematic tuning of MSAs. **a,** Correlations (Pearson’s r, abs. val.) between jaw and tongue kinematic parameters. **b,** Performance (*R*^2^) of encoding models that used the best 3 coordinate axes (jaw, tongue, or jaw+tongue) to predict smoothed spike rates, compared based on the size of the Gaussian smoothing kernel. Gray lines, single units. 25 ms std. kernel size gives the best overall performance. **c,** Parameter weights (abs. val.) for the 3 jaw, 3 tongue, and 3 jaw+tongue coordinate axes that explained the most movement variance, for all sessions. Calculated axes were generally consistent across animals/sessions, though ranked order (i.e. jaw axis 1 vs jaw axis 2) could change across animals/sessions.

**Supplemental Figure 6.**
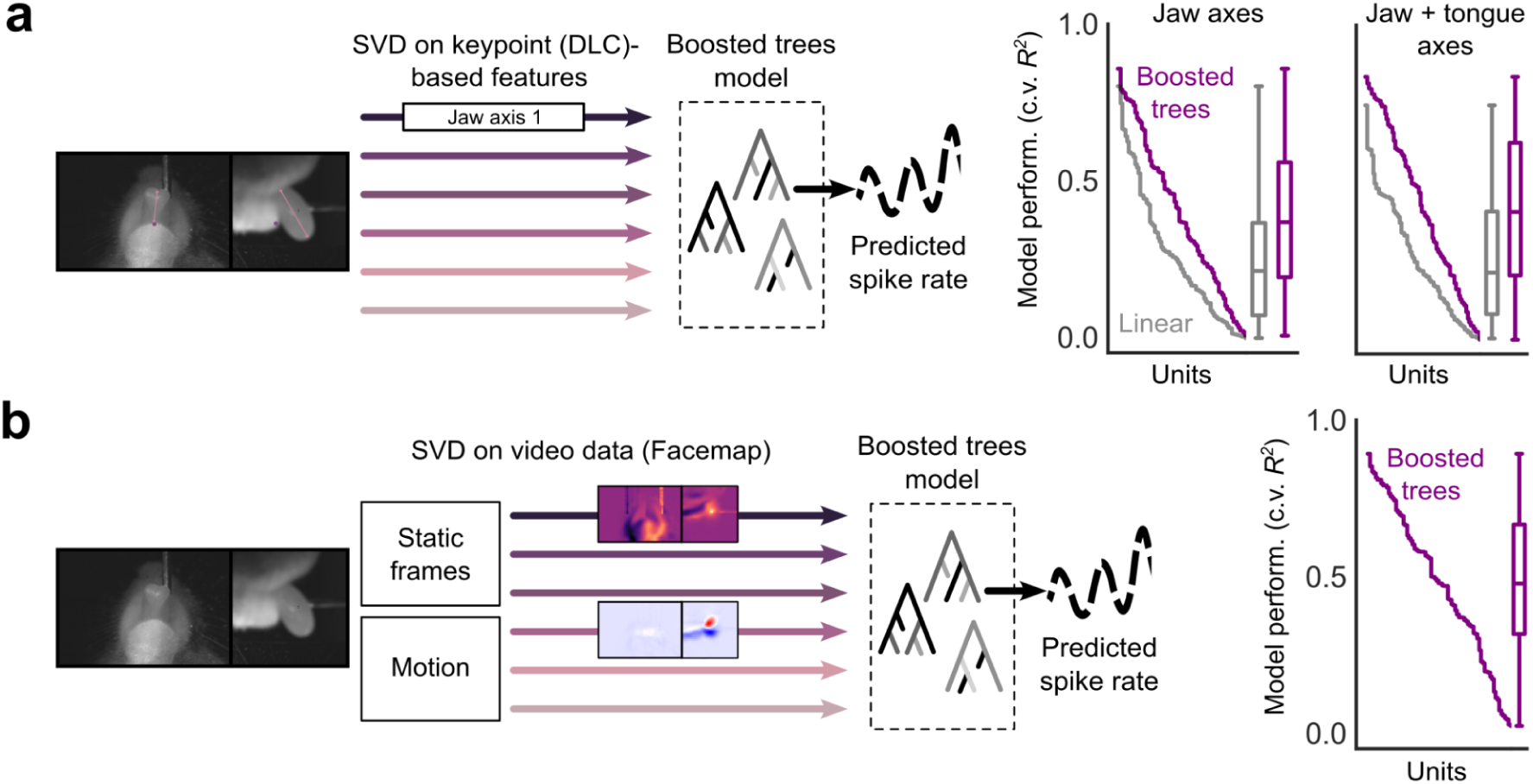
Flexible machine learning models outperformed linear models, but failed to fully explain MSA activity. **a,** Left, schematic of flexible decision-tree^16^ encoding model of unit activity. Coordinate axes extracted from tracked jaw and tongue keypoints were used to predict spike rates. Right, performance (10-fold cross-validated *R*^2^) of boosted tree models that used 6 jaw or 12 jaw+tongue axis weights. Gray indicates performance of linear models (Figure 3). **b,** Left, schematic of decision-tree encoding models after unsupervised extraction of kinematic features from the video data (Methods)^17,18^. Right, these models outperformed linear and boosted trees models that used keypoint-based data (a), however still did not explain large components of MSA activity.

**Supplemental Figure 7.**
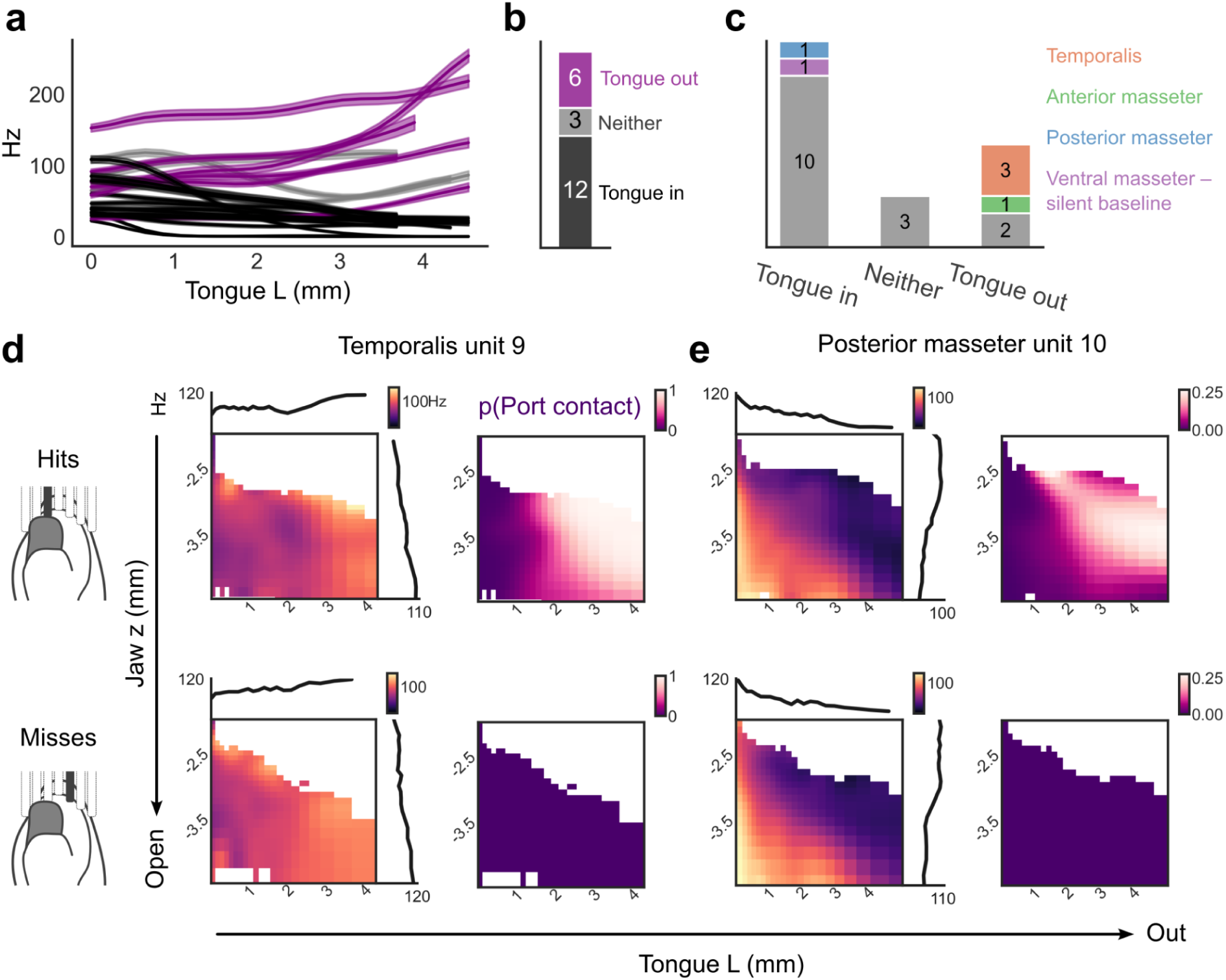
Jaw–tongue coordinate kinematic tuning was maintained in the absence of port contact. **a,** Tongue *L* tuning curves (mean ± std.) for 21 units with significant co-tuning for tongue and jaw kinematics (Figure 3n). Units were classified as ‘tongue-out’ tuned (purple) or ‘tongue-in’ tuned (black) by linear regression of firing rates to *L*. Units were tuned if p<0.05 under the null hypothesis of zero slope, with the sign of the slope indicating tuning type. **b,** Numbers of jaw*–*tongue tuning types. **c,** Cheek response field composition of units with jaw-tongue tuning. Gray indicates units with no identified cheek response field. **d*–*e,** Jaw *z*, tongue *L* 2D tuning surfaces, and 2D port contact probability surfaces, for units 9 and 10 (Figure 3), separated by ‘hit’ and ‘miss’ licks (whether the tongue contacted the port or not). While units 9 and 10 were co-tuned to orofacial conformations associated with lick port contact, co-tuning was similar in hit and miss licks.

**Supplemental Figure 8.**
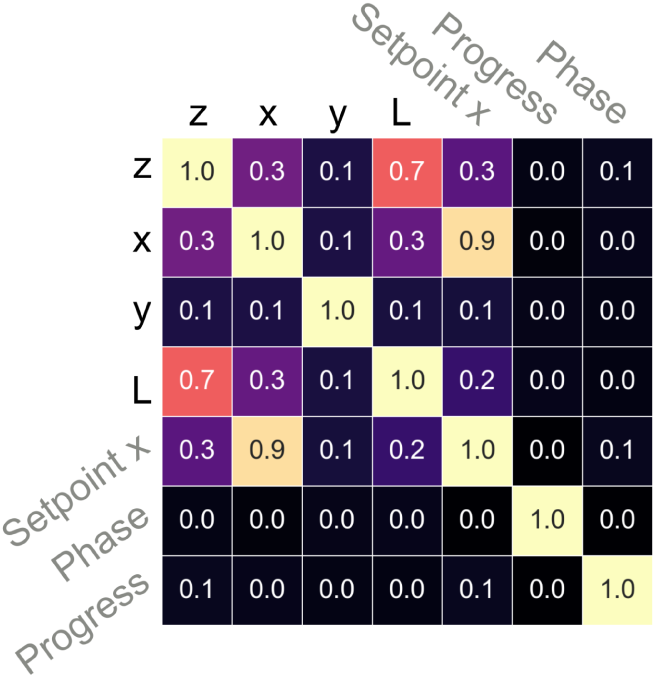
Sequence-task variables were decoupled from kinematics. Correlations (abs. val. Pearson’s r) among kinematic and task variables. While *setpoint x* was highly correlated to x, progress and phase were not linearly correlated with orofacial kinematics.

**Supplemental Figure 9.**
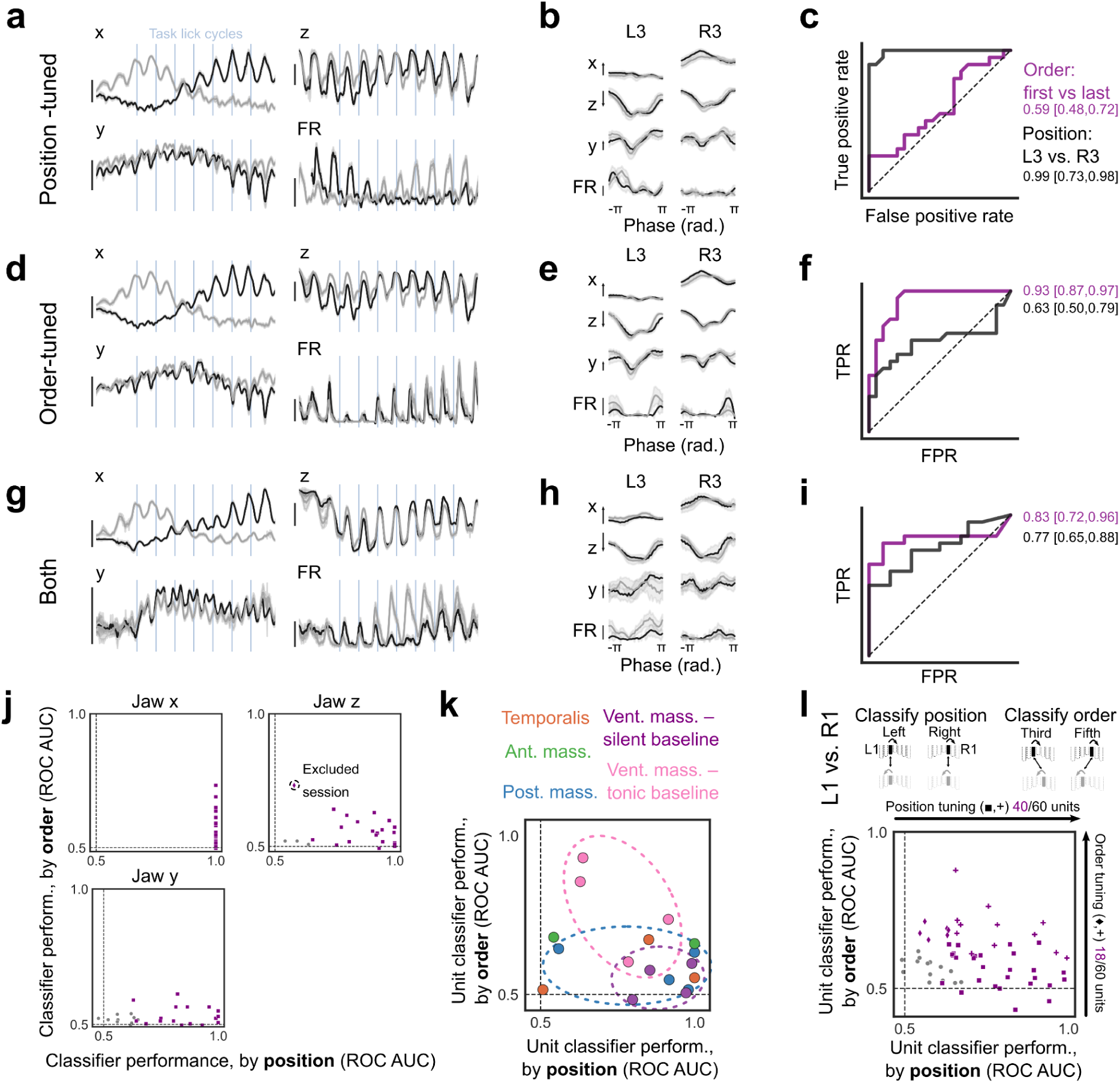
Jaw MSAs were tuned to lick position and order within sequences. **a,d,g,** Phase-aligned across-trial data for two sequences (L→R and R→L, black and gray respectively). Shown are *x*, *y*, and *z* averages (mean ± std.) and unit histograms for 3 example units (mean ± bootstrap std.). **b,e,h,** Phase-averaged lick kinematics, and phase unit histograms, for licks to L3 and R3 for the two sequences. Licks were sub-selected (n=40 total licks, 10 licks from each position-sequence combination) to match kinematics within port positions (Methods). **c,f,i,** ROC analysis for classifiers that used mean within-lick spike counts to decode port position or order. Right, AUC scores [95%CI]. c shows position tuning, f shows order tuning, and i shows tuning to both. **j,** Classifier ROC analysis that used within-lick mean jaw *x*, *z*, and *y* values for the sub-selected licks to decode position and order. Kinematics could decode lick port position but not order due to matching of licks on the kinematics. Sessions showing significant order decoding by the kinematics (dotted circles) indicated poor lick matching. Units from these sessions were excluded. **k,** Order and position classifier performance based on cheek response maps. *Posterior masseter* and *ventral masseter* – *silent baseline* units showed position tuning, while *ventral masseter* – *tonic baseline* units showed order tuning. **l,** Position and order classifier analysis that used licks to positions L1 and R1 (adjacent to the center lick). Population order tuning was weaker compared to licks to L3/R3 but was maintained in the central sequence licks. Scale bars: *z*, 1 mm open, *x*, 1 mm right, *y*, 1 mm forward, unit activity, 100 Hz.

**Supplemental Figure 10.**
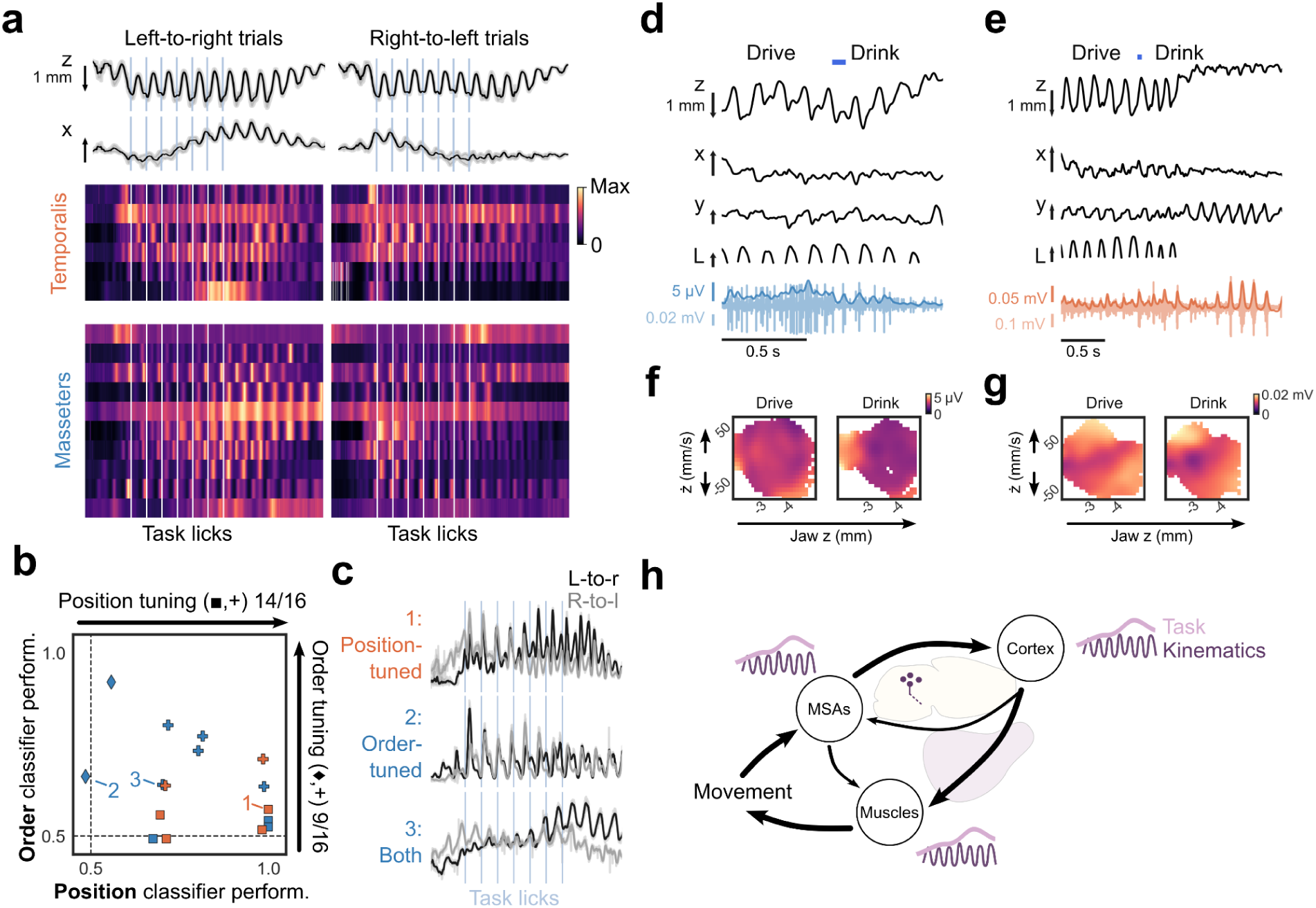
Jaw muscles show kinematic- and task-related activity during movement sequences. **a,** Phase-aligned across-trial average kinematics and rsEMG for 16 recordings. Color in heatmap is averaged rsEMG, normalized by maximum. **b**, Performance of classifiers using averaged rsEMG to classify licks to the outer two positions (L3 vs. R3) by order or position. Marker shape indicates significant classifier performance to order, position, or both. **c**, Phase-aligned across-trial average rsEMG for three recordings (indicated in f) showing position-tuning, order-tuning, or tuning to both. **d-e**, Representative jaw and tongue kinematics and rsEMG data, showing two recordings with stronger phase coupling (i.e. stronger cycle-by-cycle modulation) for drink compared to drive licks. **f-g**, Drive and drink lick 2D histograms of rsEMG for recordings in d-e averaged binned on jaw *z* and *ż*. Both recordings show sharper tuning during drink licks, indicating more complex muscle activity during drive licks. **h**, Schematic of the jaw sensorimotor system. Descending commands from cortex drive movement via the muscles and also direct top-down modulation of MSAs. MSAs flexibly encode movements, impact muscle activity via direct projections to lower motor circuits, and send ascending feedback signals back to sensorimotor cortex. Task control signals flexibly shape lower level sensorimotor transformations during motor sequences, and are therefore reflected in both feedback (MSAs) and motor output (EMG) signals.

## References

1. Merel, J., Botvinick, M. & Wayne, G. Hierarchical motor control in mammals and machines. Nat Commun 10, 5489 (2019).

2. Kleinfeld, D. et al. Low- and high-level coordination of orofacial motor actions. Current Opinion in Neurobiology 83, 102784 (2023).

3. Xu, D. et al. Cortical processing of flexible and context-dependent sensorimotor sequences. Nature 603, 464–469 (2022).

4. Scott, S. H. Optimal feedback control and the neural basis of volitional motor control. Nat Rev Neurosci 5, 532–545 (2004).

5. Todorov, E. Optimality principles in sensorimotor control. Nat Neurosci 7, 907–915 (2004).

6. Biswas, D. et al. Mode switching in organisms for solving explore-versus-exploit problems. Nat Mach Intell 5, 1285–1296 (2023).

7. Matthews, P. B. C. Muscle Spindles: Their Messages and Their Fusimotor Supply. in Comprehensive Physiology (ed. Prakash, Y. S.) 189–228 (Wiley, 1981). doi:10.1002/cphy.cp010206.

8. Blum, K. P. et al. Diverse and complex muscle spindle afferent firing properties emerge from multiscale muscle mechanics. Elife 9, e55177 (2020).

9. Ellaway, P. H., Taylor, A. & Durbaba, R. Muscle spindle and fusimotor activity in locomotion. Journal of Anatomy 227, 157–166 (2015).

10. Dimitriou, M. Human muscle spindle sensitivity reflects the balance of activity between antagonistic muscles. J Neurosci 34, 13644–13655 (2014).

11. Prochazka, A. Proprioceptive Feedback and Movement Regulation. in Comprehensive Physiology (ed. Prakash, Y. S.) 89–127 (Wiley, 1996). doi:10.1002/cphy.cp120103.

12. Prochazka, A. Sensory control of normal movement and of movement aided by neural prostheses. J Anat 227, 167–177 (2015).

13. Dimitriou, M. Human muscle spindles are wired to function as controllable signal-processing devices. eLife 11, e78091 (2022).

14. Baverstock, H., Jeffery, N. S. & Cobb, S. N. The morphology of the mouse masticatory musculature. J Anat 223, 46–60 (2013).

15. Cover, T. M. & Thomas, J. A. *Elements of Information Theory*. (Wiley, New York, 1991).

16. Chen, T. & Guestrin, C. XGBoost: A Scalable Tree Boosting System. in Proceedings of the 22nd ACM SIGKDD International Conference on Knowledge Discovery and Data Mining 785–794 (2016). doi:10.1145/2939672.2939785.

17. Syeda, A. et al. Facemap: a framework for modeling neural activity based on orofacial tracking. Nat Neurosci 27, 187–195 (2024).

18. Stringer, C. et al. Spontaneous behaviors drive multidimensional, brainwide activity. Science 364, eaav7893 (2019).

19. Warriner, C. L., Fageiry, S., Saxena, S., Costa, R. M. & Miri, A. Motor cortical influence relies on task-specific activity covariation. Cell Rep 40, 111427 (2022).

20. Bergenheim, M., Ribot-Ciscar, E. & Roll, J.-P. Proprioceptive population coding of two-dimensional limb movements in humans: I. Muscle spindle feedback during spatially oriented movements. Exp Brain Res 134, 301–310 (2000).

21. Jones, K. E., Wessberg, J. & Vallbo, A. B. Directional tuning of human forearm muscle afferents during voluntary wrist movements. J Physiol 536, 635–647 (2001).

22. Woo, S.-H. et al. Piezo2 is the principal mechanotransduction channel for proprioception. Nat Neurosci 18, 1756–1762 (2015).

23. Maxwell, D. J., Christie, W. M., Short, A. D. & Brown, A. G. Direct observations of synapses between GABA-immunoreactive boutons and muscle afferent terminals in lamina VI of the cat’s spinal cord. Brain Res 530, 215–222 (1990).

24. Fink, A. J. P. et al. Presynaptic inhibition of spinal sensory feedback ensures smooth movement. Nature 509, 43–48 (2014).

25. Koch, S. C. et al. RORβ Spinal Interneurons Gate Sensory Transmission during Locomotion to Secure a Fluid Walking Gait. Neuron 96, 1419–1431.e5 (2017).

26. Verdier, D., Lund, J. P. & Kolta, A. Synaptic Inputs to Trigeminal Primary Afferent Neurons Cause Firing and Modulate Intrinsic Oscillatory Activity. Journal of Neurophysiology 92, 2444–2455 (2004).

27. Iida, C. et al. Corticofugal direct projections to primary afferent neurons in the trigeminal mesencephalic nucleus of rats. Neuroscience 169, 1739–1757 (2010).

28. Paik, S. K. et al. γ-Aminobutyric acid-, glycine-, and glutamate-immunopositive boutons on mesencephalic trigeminal neurons that innervate jaw-closing muscle spindles in the rat: ultrastructure and development. J Comp Neurol 520, 3414–3427 (2012).

29. Kosse, C., Ivanov, J., Knight, Z., Pellegrino, K. & Friedman, J. A subcortical feeding circuit linking an interoceptive node to jaw movement. Nature (2024) doi:10.1038/s41586-024-08098-1.

30. Wood, W. W. A review of masticatory muscle function. J Prosthet Dent 57, 222–232 (1987).

31. Ribot-Ciscar, E. & Ackerley, R. Muscle proprioceptive feedback can be adapted to the behavioral and emotional context in humans. Current Opinion in Physiology 20, 46–51 (2021).

32. Papaioannou, S. & Dimitriou, M. Goal-dependent tuning of muscle spindle receptors during movement preparation. Sci Adv 7, eabe0401 (2021).

33. Clemens, A. M., Fernandez Delgado, Y., Mehlman, M. L., Mishra, P. & Brecht, M. Multisensory and Motor Representations in Rat Oral Somatosensory Cortex. Sci Rep 8, 13556 (2018).

34. Mayrhofer, J. M. et al. Distinct Contributions of Whisker Sensory Cortex and Tongue-Jaw Motor Cortex in a Goal-Directed Sensorimotor Transformation. Neuron 103, 1034–1043.e5 (2019).

35. Oryshchuk, A. et al. Distributed and specific encoding of sensory, motor, and decision information in the mouse neocortex during goal-directed behavior. Cell Reports 43, 113618 (2024).

36. Svoboda, K. & Li, N. Neural mechanisms of movement planning: motor cortex and beyond. Curr Opin Neurobiol 49, 33–41 (2018).

37. Li, N., Chen, T.-W., Guo, Z. V., Gerfen, C. R. & Svoboda, K. A motor cortex circuit for motor planning and movement. Nature 519, 51–56 (2015).

38. Bollu, T. et al. Cortex-dependent corrections as the tongue reaches for and misses targets. Nature 594, 82–87 (2021).

39. Rugh, W. J. & Shamma, J. S. Research on gain scheduling. Automatica 36, 1401–1425 (2000).

40. Azim, E. & Seki, K. Gain control in the sensorimotor system. Curr Opin Physiol 8, 177–187 (2019).

41. Confais, J., Kim, G., Tomatsu, S., Takei, T. & Seki, K. Nerve-Specific Input Modulation to Spinal Neurons during a Motor Task in the Monkey. J Neurosci 37, 2612–2626 (2017).

42. Mathis, A. et al. DeepLabCut: markerless pose estimation of user-defined body parts with deep learning. Nat Neurosci 21, 1281–1289 (2018).

43. Hill, D. N., Curtis, J. C., Moore, J. D. & Kleinfeld, D. Primary motor cortex reports efferent control of vibrissa motion on multiple timescales. Neuron 72, 344–356 (2011).

44. Chung, B. et al. Myomatrix arrays for high-definition muscle recording. eLife 12, RP88551 (2023).

45. Hill, D. N., Mehta, S. B. & Kleinfeld, D. Quality metrics to accompany spike sorting of extracellular signals. J Neurosci 31, 8699–8705 (2011).

46. Montijn, J. S. et al. A parameter-free statistical test for neuronal responsiveness. eLife 10, e71969 (2021).

47. Shigenaga, Y., Yoshida, A., Mitsuhiro, Y., Doe, K. & Suemune, S. Morphology of single mesencephalic trigeminal neurons innervating periodontal ligament of the cat. Brain Research 448, 331–338 (1988).

48. Mameli, O. et al. Role of the trigeminal mesencephalic nucleus in rat whisker pad proprioception. Behav Brain Funct 6, 69 (2010).

49. Mameli, O., Caria, M. A., Biagi, F., Zedda, M. & Farina, V. Neurons within the trigeminal mesencephalic nucleus encode for the kinematic parameters of the whisker pad macrovibrissae. Physiol Rep 5, e13206 (2017).

50. Severson, K. S. et al. Active Touch and Self-Motion Encoding by Merkel Cell-Associated Afferents. Neuron 94, 666–676.e9 (2017).

